# Disentangling mechanisms of single-cell growth rate fluctuations

**DOI:** 10.64898/2026.06.07.730294

**Authors:** Roi Holtzman, Michael Sheinman, Ariel Amir

## Abstract

Single-cell growth rates fluctuate across time and generations, but identifying the biological origin of this variability is difficult because growth rate is usually inferred from noisy measurements of cell size or mass rather than measured directly. Here, we develop a simple and interpretable framework that separates measurement noise from biological sources of growth rate variability. We show that the autocovariance of inferred instantaneous growth rates carries robust signatures of the measurement process that are largely independent of the underlying biological growth dynamics, allowing the form and magnitude of measurement noise to be identified directly from data before introducing a model for the biological dynamics. We then use the autocovariance of accumulated growth to distinguish continuous within-cycle fluctuations, division-associated perturbations, and lineage-to-lineage variability. Applying this framework to bacterial and mammalian single-cell datasets, we find evidence for continuous growth rate noise in both systems. In *E. coli*, division-associated perturbations are large at birth compared with continuous fluctuations, but their contribution to growth accumulated over the full cell cycle is reduced by rapid relaxation. In contrast, mammalian cells show no division kicks, but stronger lineage-to-lineage variability. More broadly, our results provide a direct and interpretable route to identifying the biological origin of growth rate variability in noisy single-cell measurements.

## I. INTRODUCTION

Single-cell growth rate is a central dynamical variable in cell biology. It sets the pace of biomass accumulation, couples growth to division and size control, and ultimately shapes population-level proliferation. Over the last decade, single-cell experiments have made it increasingly clear that growth is not a purely deterministic background process, but a fluctuating physiological quantity that varies across cells, across generations, and within individual cell cycles [1–7]. Understanding the source of this variability matters because it determines how single-cell fluctuations propagate to population growth [8–13] and how variability in growth contributes to broader phenotypic consequences such as developmental size heterogeneity [14] and the evolutionary effects of phenotypic variability [9, 15].

Several distinct mechanisms can underlie growth rate variability. Growth may fluctuate continuously within a cell cycle because the intracellular processes that sustain biomass production are themselves noisy [1, 4, 6]. Variability can also be introduced at division, for example through unequal partitioning of growth-related components, leading to strong division-linked perturbations, which we term “division kicks” [16]. Finally, observed growth rate differences may reflect persistent lineage-specific growth states or size-dependent growth regulation rather than ongoing stochastic fluctuations within a cycle [6, 17–2]. Distinguishing between these possibilities poses a nontrivial challenge as similar empirical behavior may reflect fundamentally different underlying mechanisms, as illustrated in Fig. 1.

**FIG. 1.**
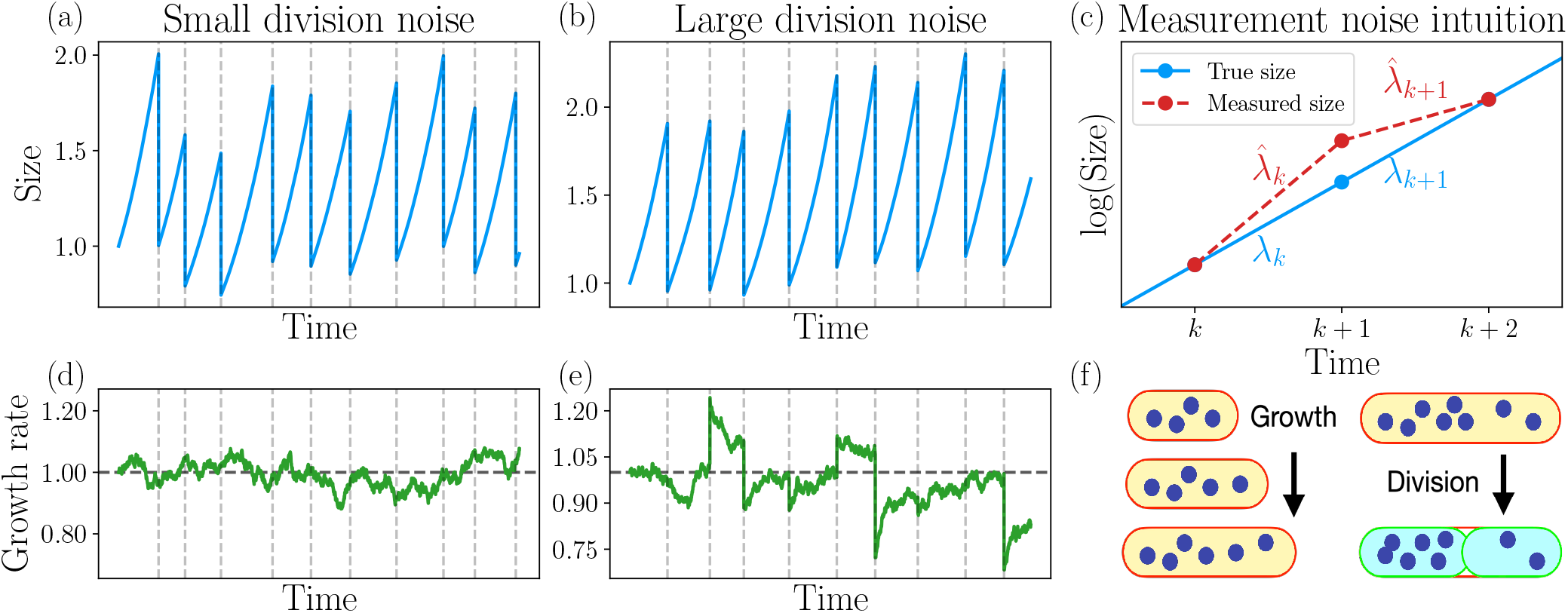
Growth rate variability. Illustration of two different mechanisms of growth rate variability and intuition for the behavior of measurement noise. Panel (f) shows plausible sources of growth rate variability. The oval shapes represent a growing cell, and the blue dots represent a protein. Such discrete evolution of proteins in low numbers, may cause continuous time fluctuations in the growth rate. At division, these proteins can partition asymmetrically, and hence cause a strong perturbation of the growth rate, which we term a “division kick”. Our goal is to distinguish between these two mechanisms in single cells experiments (Lineage-to-lineage variability is not illustrated in this plot). Panels (a,b,d,e) show this is not an easy task. Panel shows the evolution of the single-cell growth rate under continuous fluctuations (Eq. (13)), and (e) shows the evolution of additional division kicks (Eq. (15)). Vertical dashed lines mark division events. Panels (a,b) show the corresponding evolution of the cell size, driven by the growth rates in (d,e), respectively (see Eq. (12)). These show that even though the growth rate behavior is very different, the size evolution is similar, and therefore inferring the growth rate variability mechanisms is not trivial. **Measurement noise behavior**. Panel (c) illustrates the intuition behind the behavior of measurement noise. It shows the log-size of a perfect exponentially growing cell as a function of time. The blue line corresponds to the true size evolution, whereas the red line corresponds to the experimentally measured size. We consider the simple example where the sizes at times *k, k* + 2 were measured perfectly, while the size at time *k* + 1 has a measurement error. The true growth rate *λ*_*k*_ corresponds to the slope of the blue line between *k* and *k* + 1, and the measured growth rate 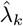 corresponds to the slope of the red line. It is clear that 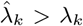. From the slopes between *k* + 1 and *k* + 2, it is clear that 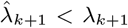, and hence measurement noise causes anti-correlations between 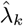 and 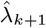.

Recent experimental advances have made these questions directly accessible. In bacteria, mother-machine microfluidic devices have enabled long-term lineage tracking over many generations under tightly controlled conditions [21]. In parallel, direct single-cell measurements of mass and volume have enabled quantitative studies of growth rate regulation across diverse organisms, from bacteria and yeast to mammalian cells [18, 20, 22–2]. Yet growth rate is usually not measured directly. Instead, it is inferred from discrete trajectories of size, mass, or volume, precisely in the regime where short-time biological changes can be comparable to measurement error. Recovering the underlying dynamics is therefore nontrivial, and several approaches have been developed to address this challenge, including statistical analyses of growth trajectories, Gaussian-process decompositions, and Bayesian inference methods [6, 28–3].

Existing inference methods can estimate model parameters once a model is specified, but they do not by themselves determine whether the assumed model captures the relevant biological mechanisms. Here we take a complementary route and develop a simple, interpretable framework that separates measurement noise from biologically distinct sources of growth rate variability as directly as possible. A central feature of our framework is that the measurement noise model can be identified separately from the biological growth model. To leading order, the signatures left by measurement noise in the auto-covariance of instantaneous growth rates, 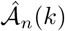,depend on the properties of the measurement errors themselves, but not on whether the underlying biological variability arises from, say, continuous fluctuations or division-associated perturbations. This makes the first step of the analysis directly interpretable: the data can be used to determine the form and magnitude of the measurement noise before introducing a model for the biological dynamics.

Having fixed the contribution of measurement noise, we then turn to the biological sources of variability using the autocovariance of accumulated growth, 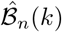. Within our framework, after properly accounting for measurement noise, the observable 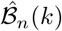 allows to distinguish the contributions of continuous fluctuations, division kicks, and lineage-to-lineage variability. We apply the method to bacterial [33] and mammalian [6] datasets. In both systems, we find evidence for continuous noise. Mammalian cells show no evidence of division kicks, but exhibit substantial lineage-to-lineage variability. In contrast, *E. coli* show strong division kicks that dominate the beginning of the cell cycle. However, they also exhibit a short relaxation time which diminishes the division kicks effect with respect to the continuous noise.

## II. DISTINCT BEHAVIOR OF MEASUREMENT NOISE

### A. Measuring growth rate from discrete size measurements

In single cell experiments, growth rate is typically inferred from discrete measurements of cell size or mass [1, 20, 23, 24]. For concreteness, we perform the analysis in terms of cell volume but the same reasoning applies to other observables such as cell mass or cell length. We denote by *v*(*t*) the true cell volume and by 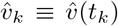 its measured value at discrete times *t*_*k*_ = *k*Δ*t*, where Δ*t* is the sampling interval and *k* is an integer. Correspondingly, *v*_*k*_ ≡ *v*(*t*_*k*_) is the true volume at time *t*_*k*_. Throughout the paper we use the hat notation to denote measured values, i.e., values that include measurement noise.

We consider additive measurement noise, namely,

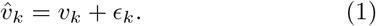

We assume that the measurement noise is Gaussian *ϵ*_*k*_ ~ 𝒩 (0, *D*_*ϵ*_), independent across measurement times 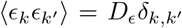, independent of the biological dynamics, and small compared to the cell volume, such that *ϵ*_*k*_*/v*_*k*_ ≪ 1.

The instantaneous growth rate is defined by

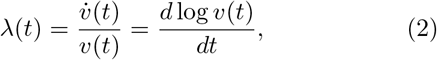

so that a constant growth rate corresponds to exponential growth in volume.

Because experiments sample cell size only at discrete times, the inferred growth rate is obtained from two successive volume measurements,

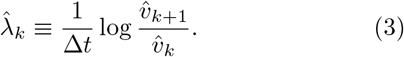

Expanding to leading order in *ϵ*_*k*_*/v*_*k*_, we obtain

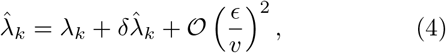

where

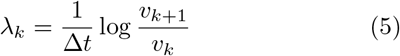

is the true discrete time growth rate, and

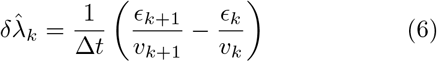

is the contribution of measurement noise to the inferred growth rate^1^.

Equation (3) for 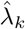 already shows why measurement noise creates a characteristic temporal structure in inferred growth rates. The same noisy measurement 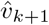 enters both 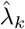 and 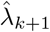 with opposite signs. As a result, measurement noise generically induces negative co-variance between adjacent inferred growth rates. Finally, we note that a centered finite-difference definition for 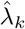 does not change our results qualitatively as the same overall framework and conclusions remain the same, see Supplementary Information (SI) V A.

### B. Autocovariance of the measured growth rate

Autocovariance of the inferred growth rate provides a natural way to quantify temporal structure in growth rate fluctuations. Since the inferred growth rate contains both biological variability and measurement error, the first step is to determine how measurement noise contributes to this autocovariance.

To this end, we define the dimensionless autocovariance of the instantaneous growth rates at time index *k* and lag *n*,

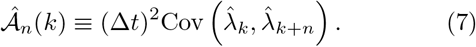

The prefactor (Δ*t*)^2^ makes 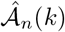 dimensionless and removes the trivial scaling with the sampling interval.

Using the expansion in Eq. (4), and the assumption that the measurement noise is independent of the biological dynamics, we obtain

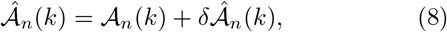

where 𝒜_*n*_(*k*) ≡ (Δ*t*)^2^Cov (*λ*_*k*_, *λ*_*k*+*n*_) is the contribution of the true growth rate dynamics, and 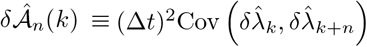 is the measurement noise contribution.

At this stage, no specific model for *λ*(*t*) is needed. We first analyze 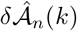, the structure of which is determined entirely by the way measurement noise enters the inferred growth rate. Only afterwards do we introduce a biological model and calculate 𝒜_*n*_(*k*).

### C. Intuition for the measurement noise contribution to 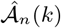

The measurement noise contribution to 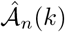 has a simple and universal structure across lags *n*. It is positive at lag *n* = 0, negative at lag *n* = 1, and vanishes for *n* ≥ 2. This structure follows directly from the way measurement noise enters the inferred growth rate in Eq. (3). To see the origin of the negative autocovariance at lag *n* = 1, consider the toy example shown in Fig. 1, where the measured value 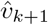 lies above the true trajectory, while the values at *t*_*k*_ and *t*_*k*+2_ are measured without error. In this case, the inferred growth rate between *t*_*k*_ and *t*_*k*+1_ is overestimated, 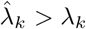, because the segment connecting 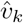 and 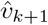 is steeper than the true one. At the same time, the inferred growth rate between *t*_*k*+1_ and *t*_*k*+2_ is underestimated, 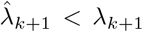, because the segment connecting 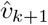 and 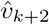 is less steep than the true one. Thus, the same positive measurement error in the volume at time *t*_*k*+1_ drives 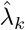 upward and 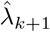 downward, producing negative covariance between adjacent growth rates. Naturally, the same reasoning applies when the measurement error has the opposite sign. This behavior is somewhat reminiscent of “regression toward the mean” [34]. Here, however, it is not the true growth rate that regresses toward the mean, but the estimate inferred from size measurements.

By contrast, for *n* = 0, measurement noise contributes positively to the variance of the inferred growth rate. For *n* ≥ 2, the inferred growth rates 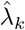 and 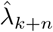 do not share any common measurement error term. Therefore, under the assumption of independent measurement noise, their covariance vanishes.

This qualitative argument already shows that measurement noise generates a characteristic lag dependence in 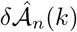. In the next subsection, we derive the corresponding expression for 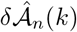 explicitly.

### D. Measurement noise signatures

The contribution of measurement noise to 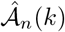 can be calculated explicitly and its full form is given in SI V. Here, we present a simpler expression which is obtained by assuming that cells grow exponentially on average, and that fluctuations of *λ*(*t*) around the mean growth rate *λ*_0_ are small. We further assume *λ*_0_Δ*t* ≪ 1, which holds when each cell cycle is sampled at many time points. Under these assumptions, substituting 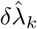, (6), into Eq. (8), we find

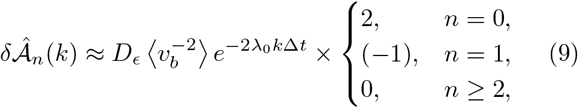

where *v*_*b*_ is the cell volume at birth. Since the measurement noise is additive, its relative effect decreases as the cell grows, and therefore 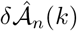 decreases with *k*.

Equation (9) yields two simple and robust signatures. First, the contributions at lags 0 and 1 obey

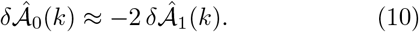

Second, because the cell approximately doubles its size over one cell cycle, we have 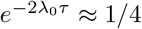, where *τ* is the division time, and ⟨*τ* ⟩ is its mean. Therefore

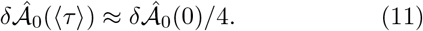

To test the predictions in Eqs. (9), (10) and (11), we analyzed the experimental data obtained from length measurements of *E. coli* cells grown in a mother-machine apparatus [33] and mass measurements of mammalian mouse lymphocytic leukemia L1210 cells using suspended microchannel resonators [6]. In Fig. 2, we plot 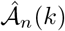 for both datasets (further *E. coli* datasets are plotted in SI XIII^2^). As Eq. (9) predicts, we observe that 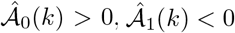, and 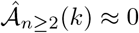. In addition, 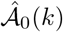 approximately overlaps with 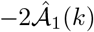 (recall Eq. (10)), and its value near the mean division time is close to one quarter of its initial value (recall Eq. (11)). Taken together, these observations strongly suggest that the observed autocovariance is dominated by measurement noise. This consistency supports the assumptions that the measurement noise is additive and independent. Indeed, if the noise were multiplicative, 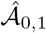 would not show the observed decrease with *k*; if it were not independent, the factor of 2 in (10) would change, and, in general, 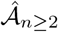 would be nonzero. See SI IX and SI VIII for further details on these extensions.

**FIG. 2.**
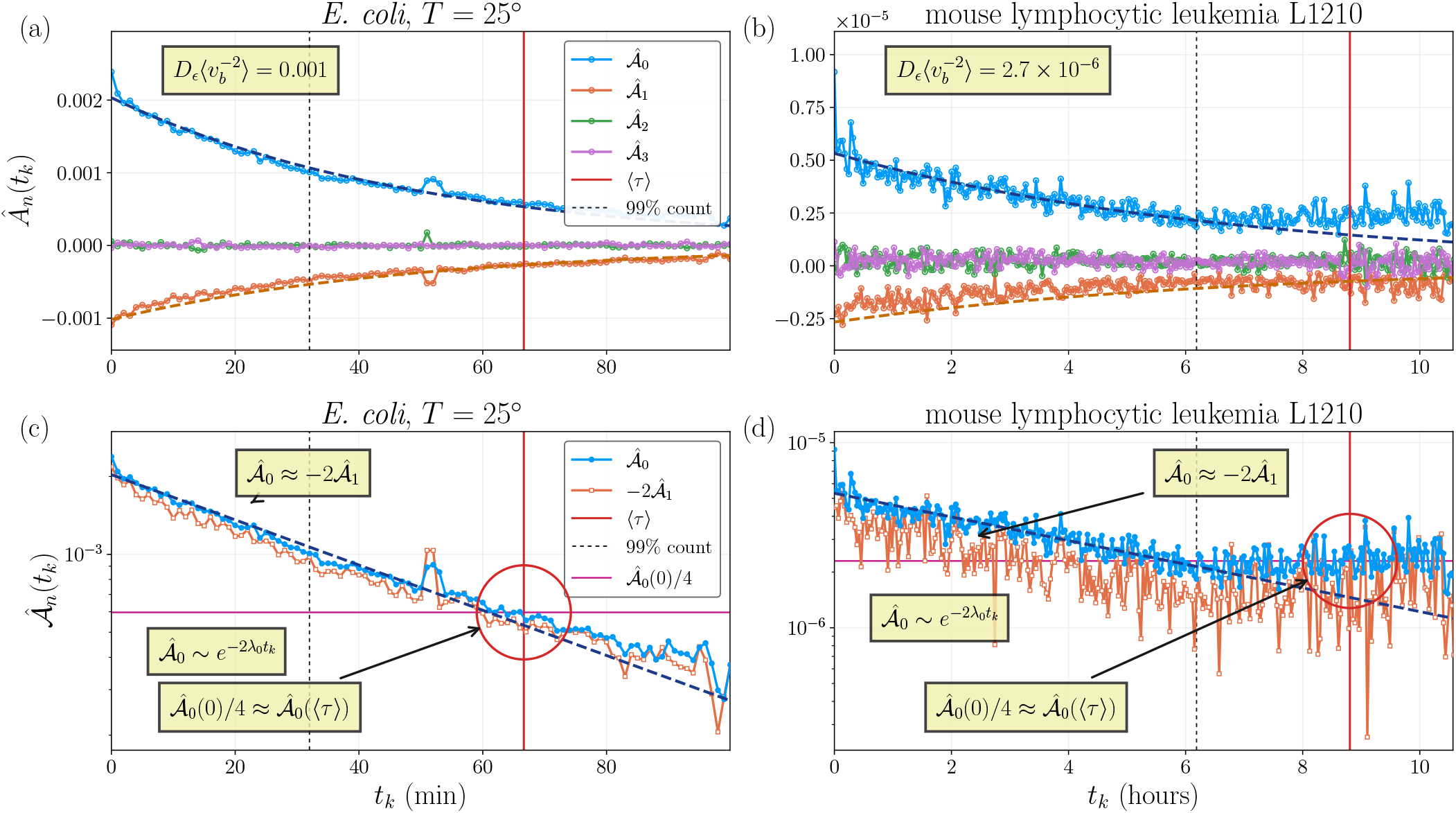
Instantaneous growth rate autocovariance 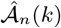 for different lags *n* for *E. coli* [33], and mammalian cells [6]. Analyses of *E. coli* experiments at different temperatures are shown in Fig. 3 in SI XIII. The top panels show that 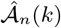 follows the simple behavior predicted in Eq. (9): 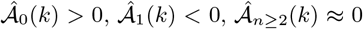. The dashed dark blue curves are exponential fits, 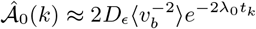, which provide an estimate for the measurement noise magnitude. The orange dashed line in the top panel corresponds to 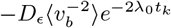, using the parameters from the fit of 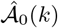. The bottom panels show 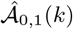 in semi log-scale, showcasing the exponential behavior. Moreover, they show the predictions 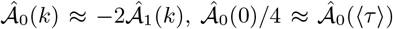 from Eqs. (10), (11), respectively. The red line is the mean division time ⟨*τ*⟩ obtained from the data. The black vertical dashed line marks the moment in time that the cell count is 99% from the initial count.

Our analysis was carried out in absolute cell age. As a result, the statistics 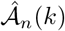 is calculated using a set of cells that changes with time, since it contains only cells that have not yet divided by age *t*_*k*+*n*+1_. The effects of this survival bias can be taken into account within a specified growth and division model, and are discussed in SI VII. In practice, restricting the analysis to times with high survival fraction mitigates this bias.

Having characterized the properties of the measurement noise in 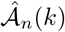, we next introduce a model for the true growth dynamics and derive the corresponding form of 𝒜_*n*_(*k*).

## III. MODEL FOR BIOLOGICAL GROWTH

### A. Cell cycle, growth, and division

We model growth at the level of a single cell cycle. A cell is born at time *t* = 0 with volume *v*_*b*_ ≡ *v*(0) and initial growth rate *λ*_*b*_ ≡ *λ*(0). The central dynamical variable in our description is the instantaneous growth rate *λ*(*t*), which determines the volume through

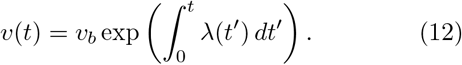

The cell cycle ends at division time *τ*, when the cell reaches a division volume *v*_*d*_ = *v*(*τ*). To allow variability in division timing, we assume a noisy size control rule, which determines *v*_*d*_ from the birth volume *v*_*b*_ together with an additional stochastic contribution. This family of size regulation models interpolates between timers, adders and sizers via a parameter 0 *< α <* 2 [35–37] (see more details in SI I). The behavior of the growth rate statistics is not sensitive to *α*, but in order to compare simulations and experiments, we used values fitted from the data. The division time *τ* is determined by solving Eq. (12), setting *v*(*τ*) = *v*_*d*_.

At division, the mother cell divides to two daughter cells, and time is reset so that each daughter begins a new cycle at *t* = 0. In the absence of division kicks, the growth state is inherited continuously across division, so that the daughter starts with the mother’s final growth rate. Below we generalize this by allowing an additional stochastic perturbation to the growth rate at birth.

### B. Stochastic growth rate

We model biological variability in the growth rate through three contributions: continuous fluctuations during the cell cycle, possible perturbations at birth, and lineage-to-lineage variability in the mean growth state.

#### 1. Continuous fluctuations during the cell cycle

Within our model, during a cell cycle, the growth rate follows an Ornstein-Uhlenbeck (OU) process [12, 38, 39],

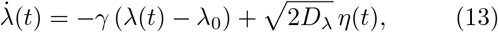

where *λ*_0_ is the mean growth rate, *γ* = 1*/τ*_rel_ is the relaxation rate, *D*_*λ*_ sets the strength of the fluctuations, and *η*(*t*) is Gaussian white noise with

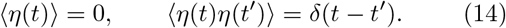

This provides a simple model of continuous intracellular fluctuations around a preferred growth state. The key timescale is the relaxation time *τ*_rel_, which will later be compared with the cell cycle duration *τ* ≈ log 2*/λ*_0_.

#### 2. Division kicks

To allow for perturbations generated at cell division, we introduce division kicks. These represent birth associated shifts in the growth rate, for example due to asymmetric partitioning of intracellular components at division [16]. We model them as an additive Gaussian perturbation to the inherited growth rate,

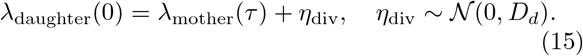

Thus, division kicks act only at birth, after which their effect relaxes during the cell cycle through the same continuous dynamics described above. A schematic example of growth trajectories with and without division kicks is shown in Fig. 1.

#### 3. Lineage variability

In addition to stochastic fluctuations within a cell cycle, different lineages may have different typical growth rates [6, 17, 40]. To account for this, we let the mean growth rate depend on the lineage,

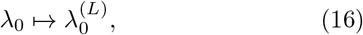

where *L* labels the lineage. We model the lineage dependent mean 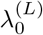 as a random variable drawn once per lineage from a Gaussian distribution with mean *λ*_0_ and variance *D*_ℓ_.

Ideally, one would estimate growth statistics separately for each lineage. In practice, however, experimental datasets often do not contain enough observations per lineage for reliable estimation. We therefore analyze pooled statistics across lineages, and treat lineage variability as an additional source of biological variation at the population level.

## IV. MEASUREMENT NOISE DOMINATES 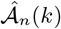

We now turn to the biological contribution 𝒜_*n*_(*k*) to the autocovariance of the inferred growth rate. Within the model introduced above, 𝒜_*n*_(*k*) can be written as

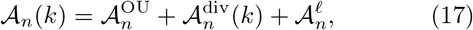

where 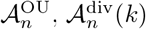, and 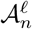 denote the contributions of the OU fluctuations, division kicks, and lineage variability, respectively. Closed form expressions for these terms are derived in SI III.

The full expressions depend on the relation between the relaxation time 1*/γ* and the cell cycle duration. Here, we focus on the regime *γ*⟨ *τ*⟩ ≫ 1 in which relaxation is fast compared with the mean cell cycle duration, so that memory of the growth rate at birth is lost well before division, see e.g. [1]. This regime yields simple analytical expressions and is also the most relevant for the fitted datasets below. The more general case, including *γ* ⟨*τ*⟩ ≲ 1, is discussed in SI II. As we show there, the main conclusion of this section remains unchanged: the biological contribution to 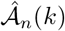 does not reproduce the characteristic lag structure generated by measurement noise.

In the fast relaxation regime, *γ*⟨*τ* ⟩ ≫ 1, we obtain

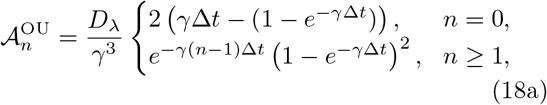

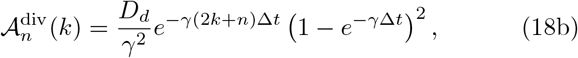

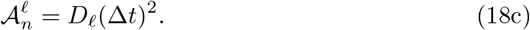

These expressions show that all three biological contributions are nonnegative. The OU term is stationary in *k* and decays with lag *n*. The division kick term decays with both lag *n* and age *k*. The lineage variability term is independent of both *k* and *n*.

This structure is qualitatively different from the measurement noise contribution, Eq. (9). In particular, the negative lag *n* = 1 autocovariance observed in the data, both in *E. coli* [33] and mammalian cells [6], cannot be explained by the biological mechanisms considered here, and is instead a distinctive signature of measurement noise. We therefore conclude that in these datasets the observed autocovariance is dominated by the measurement noise contribution, 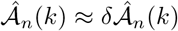. Furthermore, the form in Eq. (9), and its characteristic signatures (10), and (11), agree very well with both datasets. This conclusion allows us to estimate the measurement noise magnitude from the fit of 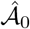 in Fig. 2. For the mammalian dataset we obtain 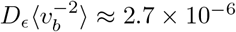. This is in agreement with Ref. [6] which reports mass measurement error of ~ 0.1%, which in our notation corresponds to 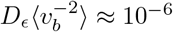

The dominance of measurement noise in 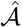 can also be understood from a simple scaling argument. Inferring instantaneous growth rates requires small sampling interval Δ*t*. In the experimentally relevant regime *γ*Δ*t* ≪ 1, Eqs. (17) and (18) give

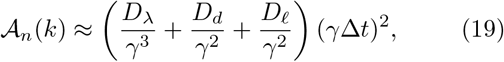

whereas the measurement noise contribution in Eq. (9) does not scale with Δ*t* in this way. Thus, improving temporal resolution suppresses the biological contribution to 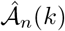 relative to the measurement noise contribution. This explains why 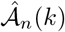 is particularly well suited for estimating both the features (i.e., additive and independent) and the magnitude of measurement noise, but is generally less informative about the underlying biological fluctuations. This suggests that the measurement noise dominated behavior seen in the experiments arises from the relative scaling of the two contributions, rather than from an absence of biological fluctuations.

## V. ACCUMULATED GROWTH COVARIANCE REVEALS BIOLOGICAL NOISE MECHANISMS

Since 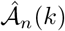 is generically dominated by measurement noise, we seek an observable that highlights the biological signal. We therefore, similarly to Refs. [6, 12], consider the accumulated growth since birth up to time *t*_*k*_, defined by

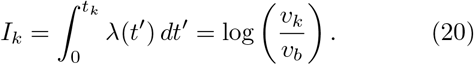

The corresponding measured quantity is

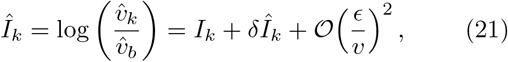

where

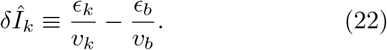

In analogy with 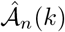, we define the autocovariance of the measured accumulated growth by

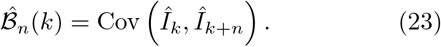

Since 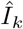 is dimensionless, 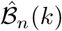 is also dimensionless.

Using Eq. (21), and the independence of measurement noise from the biological dynamics, 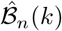 decomposes as

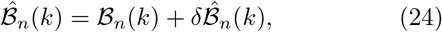

where ℬ_*n*_(*k*) ≡ Cov (*I*_*k*_, *I*_*k*+*n*_) is the biological contribution 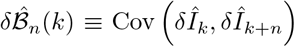 is the contribution of measurement noise.

The measurement noise contribution is derived in SI V and is given by

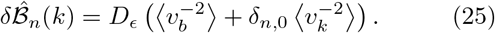

The biological contribution again decomposes (compare with Eq. (17)) into the three mechanisms introduced above^3^,

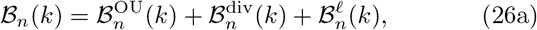

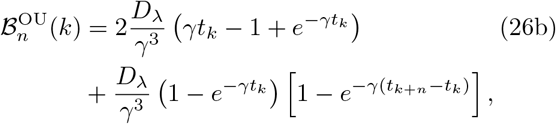

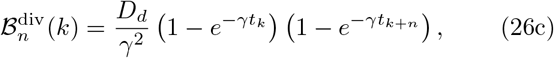

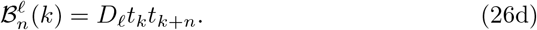

In contrast to 𝒜_*n*_(*k*), the terms in ℬ_*n*_(*k*) grow with age and are therefore not suppressed relative to measurement noise. This makes 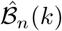 useful for distinguishing biological mechanisms.

The form of Eq. (25) suggests that it is particularly convenient to focus on lag *n* = 1. For all *n* ≥ 1, the measurement noise contribution is the same constant 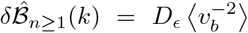, while the biological terms are largest and simplest at small lag. Therefore, we analyze 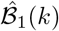.

For *n* = 1, and assuming *γ*Δ*t* ≪ 1, we find the following forms for early *t*_*k*_ ≪ 1*/γ* and late *t*_*k*_ ≫ 1*/γ* times

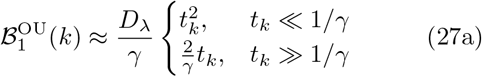

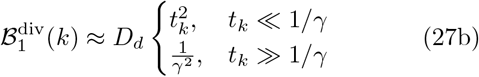

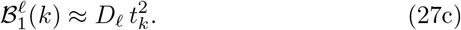

Thus, at early times, all three biological contributions have the same leading quadratic dependence and cannot be distinguished. In contrast, at late times, continuous fluctuations produce a contribution that grows linearly with age, division kicks produce a plateau, and lineage variability produces a quadratic growth. These different trends corresponding to different biological noise mechanisms allow for qualitatively testing the compatibility of the model’s predictions against experimental data.

For 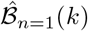, the measurement noise contribution is a constant 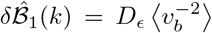, and is therefore indistinguishable from the division kicks contribution 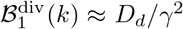. Usefully, 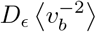 is estimated independently from the fit of 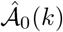 to Eq. (9). This then allows us to subtract the measurement noise contribution 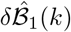 from 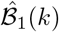 using Eq. (25). We therefore define the shifted accumulated growth autocovariance

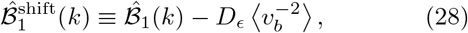

and fit it to the biological contributions ℬ_1_(*k*) in Eqs. (26).

The relaxation time scale *τ*_rel_ = 1*/γ* largely affects the behavior of the biological noise mechanisms 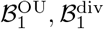 with time, see Eqs. (26). The other parameters, *D*_*λ*_, *D*_*d*_, *D*_ℓ_, only determine the magnitudes of these different noise mechanisms. As there is no *a priori* way to determine the value of 1*/γ*, we scan over a broad range of its plausible values, and fit the other parameters *D*_*λ*_, *D*_*d*_, *D*_ℓ_ for each value of 1*/γ* using a standard mean-squared-error of 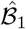, see Eq. (158) in the SI. We find that the loss has a global minimum as a function of 1*/γ*, with a smooth neighborhood around it, see Fig. 1 in SI XII B.

Using these fitted values, we then simulated the model and produced similar datasets faithful to the experiments. Namely, each simulation has the same number of lineages, and the same number of cycles within each lineage. Then, performing 1000 realizations per experimental dataset using the parameters obtained from the experimental fits, we follow the same fitting procedure to obtain a set of fitted parameters per simulation. From these simulations we obtain confidence intervals for our model given the experimental data. Values and histograms of the fitted parameters are given in Table I and plotted in Fig. 2 in SI XII.

Before analyzing the resulting fits, it is useful to clarify how we compare the magnitude of fluctuations generated by division kicks with those generated by continuous noise. Division kicks act as impulsive perturbations at birth and then relax during the cell cycle, whereas continuous fluctuations are present throughout the cycle. A natural first comparison is the ratio of the perturbation magnitudes at birth, *R*_birth_ = *D*_*d*_*/*(*D*_*λ*_*/γ*). However, this ratio does not quantify the total effect of division kicks over the cell cycle. To capture this effect, we compare the contributions of the two sources to the variance of accumulated growth over one typical cycle, as measured by ℬ_0_:

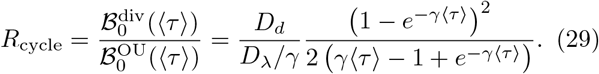

In the fast-relaxation limit, *γ*⟨*τ* ⟩ ≫ 1, this becomes *R*_cycle_ ≈ *R*_birth_*/*(2*γ*⟨*τ* ⟩) ≪ *R*_birth_. Thus, even when *R*_birth_ ≫ 1, so that division kicks dominate the perturbation immediately after birth, their contribution to accumulated growth can be strongly suppressed when the relaxation time *τ*_rel_ = 1*/γ* is short compared with the mean cell-cycle duration.

We next analyze the fits shown in Fig. 3. For the *E. coli* dataset, the shifted autocovariance 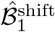 is best described by a short relaxation time, *τ*_rel_*/* ⟨*τ*⟩ ≈ 0.03. Division kicks dominate the instantaneous variance at birth, with *R*_birth_ ≈ 13. However, because they relax rapidly, their contribution to growth accumulated over the full cell cycle is much smaller, *R*_cycle_ ≈ 0.22. This interpretation is consistent with Fig. 3: the fitted division kick contribution is largest near birth and decays rapidly, while the later time autocovariance is dominated by the OU and lineage components. Additional *E. coli* datasets are shown in SI XIII and provide similar values of *R*_birth_ and *R*_cycle_. For the mammalian dataset, we find only continuous OU noise and no evidence for division kicks. Moreover, the late-time behavior is captured by lineage variability together with a relaxation time of *τ*_rel_*/* ⟨*τ*⟩ *≈* 0.22. We note that Ref. [6] also found no evidence for division kicks using a Gaussian-process framework and maximum-likelihood estimation, although they estimated a longer relaxation time, *τ*_rel_*/* ⟨*τ*⟩ *≈* 0.53, which is within our confidence interval; see Table I in SI XII.

**FIG. 3.**
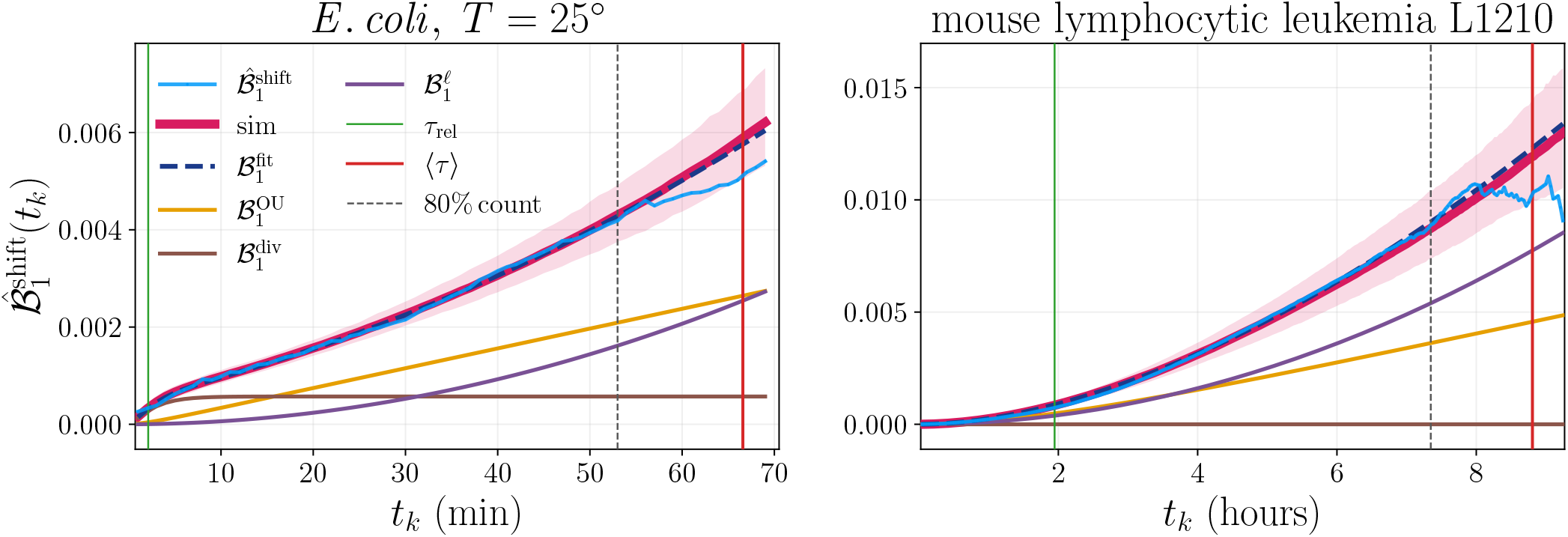
The shifted accumulated growth rate autocovariance 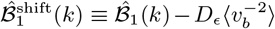 for *E. coli* [33], and mammalian cells [6]. Analyses of *E. coli* experiments at different temperatures are shown in Fig. 4 in SI XIII. The constant 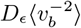 is obtained from the fit in Fig. 2. The light blue line is the data, and the dark blue dashed lines are fits of 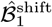 using Eq. (26), performed as described in SI XII B. From the fit, we plot separately the different contributions: OU noise (yellow), division kicks (brown), and lineage variability (purple), see Eq. (27). In pink, we plot the median of 1000 realizations of simulations performed with the values of *γ, D*_*λ*_, *D*_*d*_, *D*_*ϵ*_, *D*_ℓ_ obtained from the experiments, as described in SI XII C, XII D. The shaded pink ribbon marks the 95% confidence interval, see Table I in SI XII. The green line denotes the fitted relaxation time *τ*_rel_ = 1*/γ*. The red line is the mean division time ⟨*τ*⟩ obtained from the data. The vertical black dashed line marks the threshold for the data we used for the fit. We set this threshold to 80% count of the initial number of cells in order to mitigate the effects of survival bias.

We conclude this section by noting that measurement noise and division kicks can generate similar signatures in the data. Therefore, to estimate the magnitude of division noise reliably, one must first estimate the measurement noise and properly remove its contribution.

## VI. GENERALIZATIONS OF MEASUREMENT NOISE MODEL AND GROWTH LAW

In our analysis we assumed Gaussian, additive, and independent measurement noise, together with the growth rate definition in Eq. (2), which is natural for exponential growth. Next, we examine how the autocovariances signatures change when these assumptions are relaxed.

### A. Correlated measurement noise

The general case is discussed in SI VIII. Here, we only show the results for the simple case of exponentially correlated measurement noise, ⟨*ϵ*_*k*_*ϵ*_*k*+*n*_⟩ = *D*_*ϵ*_*ρ*^|*n*|^ with |*ρ*| *<* 1.

In this case, the measurement noise contribution to 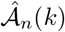 is no longer strictly local in lag *n*. To leading order, we obtain

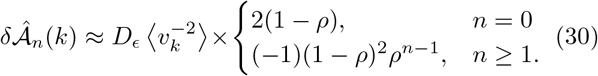

Thus, temporal correlations in the measurement noise broaden the lag structure of 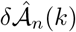 beyond the uncorrelated case in Eq. (9). We note that the relation between the lags *n* = 0, 1 in Eq. (10) gets an extra factor of (1 − *ρ*)^−1^, such that 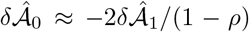. Moreover, 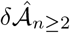 is no longer zero, but diminishes exponentially with lag *n*. The sign of 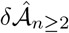 depends on the sign of *ρ*. For 0 *< ρ <* 1, the contribution at lags *n*≥ 2 remains negative, whereas for −1 *< ρ <* 0, its sign alternates with lag. Therefore, correlated measurement noise preserves the basic idea that the growth rate estimator itself induces a structured autocovariance pattern, but the pattern is no longer restricted to the first two lags. This analysis further supports our assumptions of uncorrelated measurement noise.

### B. Multiplicative measurement noise

We also consider multiplicative measurement noise. In that case, the relative magnitude of the measurement noise does not decrease as the cell grows. As a result, 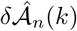 does not decay with age. This behavior is qualitatively different from the experimental datasets analyzed in Fig. 2, where the magnitude of 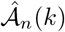 decreases over the cell cycle. This supports the additive noise assumption used in the analysis. See details in SI IX.

### C. Linear growth

Finally, we consider cells that grow linearly in size rather than exponentially [30, 41]. In this case, it is natural to redefine the growth rate as

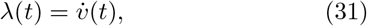

rather than 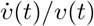 as in Eq. (2). With additive measurement noise, we find that the corresponding autocovariance of the inferred growth rate does not decay with age. This is similar to the behavior obtained for multiplicative noise under exponential growth. More details and explicit formulas are in SI X.

## VII. DISCUSSION

In this work, we developed a two-step framework for analyzing noisy single-cell growth data. We showed that when instantaneous growth rates are inferred from adjacent size measurements, measurement noise leaves simple and robust signatures in the growth rate autocovariance statistics, denoted by 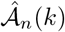. This statistic can be decomposed into a biological contribution and a measurement noise contribution, 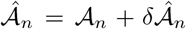. We further argued, and verified in two experimental datasets, that in practice the measurement noise term is expected to dominate. Intuitively, increasing the temporal resolution, corresponding to smaller Δ*t*, suppresses the biological contribution 𝒜_*n*_, while leaving the measurement noise contribution 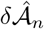 of the same order. As a result, 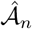 acquires a characteristic pattern: it is positive at lag *n* = 0, negative at lag *n* = 1, and zero for all lags *n* ≥ 2. The negative lag-one covariance arises naturally because the same noisy size measurement appears in the estimators of two consecutive growth rates, thereby inducing anticorrelations between them. Since this effect is determined entirely by the measurement procedure, it is largely independent of the biological source of noise. This makes the autocovariance signature a useful diagnostic for estimating the magnitude of measurement noise and for probing whether its assumed form, e.g. additive and independent, is consistent with the data.

We then introduced the covariance of accumulated growth, 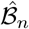, for which the biological contribution grows with cell age whereas the measurement noise contribution remains constant. Using the measurement noise amplitude inferred from 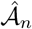, we fitted 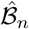 to a biological growth model. This analysis shows that the *E. coli* data exhibit strong division kicks at birth, *R*_birth_ ≈ 13. However, because these kicks relax rapidly, their contribution to accumulated growth over the full cell cycle is reduced to *R*_cycle_ ≈ 0.22. By contrast, the mammalian data show no evidence for division kicks, and are instead described by continuous growth rate fluctuations together with strong lineage-to-lineage variability.

More generally, the logic of the framework is not tied to this specific model choice. Once measurement noise has been identified and quantified through 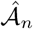, the observable 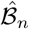 can be analyzed within other biological models of growth fluctuations as well. In our case, different biological noises contributed additively to 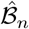 in Eq. (26), making it easy to add and remove noise mechanisms from the model.

Our work relates to that of Kiviet *et al*. [1], where the instantaneous growth rate was obtained by fitting local exponentials to the cell length trajectory. In that sense, their reported “instantaneous” growth rate is already a locally fitted, and therefore locally smoothed estimator. Our framework instead addresses the complementary setting in which growth rate is inferred directly from discrete size measurements.

More recent approaches use explicit principled inference procedures. Levien *et al*. [6] infer growth fluctuations of mammalian cells using Gaussian-process smoothing and detrending. They conclude that within-lineage growth variability is dominated by continuous growth fluctuations rather than by division kicks, which is consistent with our result. A recent Bayesian approach [29] performs full model-based inference by maximizing the likelihood of the noisy measurements and reconstructing posterior cell-state trajectories with uncertainty estimates. By contrast, our method is formulated directly in terms of covariance observables inferred from discrete size measurements. This makes the role of measurement noise explicit and yields a more interpretable separation between measurement noise, continuous growth fluctuations, and division kicks.

A related point is that our observables are covariances, and therefore are insensitive to a deterministic absolute time trend in the growth rate, provided that this trend is common across cycles and is accounted for in the mean. By an absolute time trend we mean a deterministic function *λ*_trend_(*t*) that replaces the constant mean growth rate *λ*_0_. This extension is discussed in SI XI.

In conclusion, empirical estimation of instantaneous growth rate fluctuations is challenging because measurement noise can obscure the short time biological signal. Smoothing-based approaches improve robustness, but they also average the dynamics over longer times. Statistical inference procedures are powerful for estimation, but their mechanistic interpretation can be less direct. The present work provides an alternative route: it uses the covariance signature generated by measurement noise to identify and quantify that noise, and then exploits a second observable to distinguish biologically meaningful fluctuation mechanisms. In this sense, the main value of the framework is not only parameter estimation, but also transparent assessment of whether the assumed noise and growth models are consistent with the data.

## Acknowledgments

It is a pleasure to thank Naama Brenner, Kuheli Biswas, Ethan Levien, Teemu Miettinen, and Jaan Mannik for fruitful discussions. A.A. acknowledges support from the European Union (ERC, BIGR, 101125981), the Israeli Science Foundation (146873) and the Clore Center for Biological Physics. R.H. acknowledges support from the Leverhulme Trust International Professorship Grant (No. LIP-2020-014) at the University of Oxford, and the Faculty Postdoctoral Excellence Fellowship of the Weizmann Institute of Science.

## Footnotes

1 We note that even the true growth rate *λ*_*k*_ (i.e. when measurement noise is zero) is not instantaneous but rather is the average of the growth rate over an interval 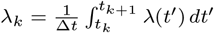.

2 In the main text, for *E. coli*, we included data corresponding to *T* = 25° since the cell cycle is longer and has more data points. The temperature *T* = 37° is in SI XIII.

3 Here we consider the fast relaxation regime *γ* ⟨*τ*⟩ ≫ 1 corresponding to Eq. (18). The general case is given in SI IV.

