## Supplementary Material for "Disentangling mechanisms of single-cell growth rate fluctuations"

### Supplementary Material for
### Disentangling mechanisms of single-cell growth rate fluctuations

Roi Holtzman*
*Rudolf Peierls Centre for Theoretical Physics,*
*Clarendon Laboratory, Oxford OX1 3PU, UK and*
*Department of Physics of Complex Systems,*
*Weizmann Institute of Science, Rehovot, 76100, Israel*

Michael Sheinman† and Ariel Amir‡
*Department of Physics of Complex Systems,*
*Weizmann Institute of Science, Rehovot, 76100, Israel*
(Dated: June 5, 2026)

#### CONTENTS

I. Model and definitions 2

II. Continuous-time autocovariance of the biological growth rate 3

III. Statistics of the instantaneous growth rate 4

IV. Statistics of the accumulated growth rate 5

V. Measurement noise 6
A. Alternative centered finite-difference estimator 8

VI. Asymptotic behavior of $\mathcal{B}_1(k)$ 9

VII. Survival bias in age-conditioned statistics 9
A. Applications to $\mathcal{A}_n(k), \mathcal{B}_n(k)$ 12

VIII. Correlated measurement noise 13

IX. Multiplicative measurement noise 14

X. Linear growth 15

XI. Deterministic trend in the growth rate 16

XII. Data analysis from experiments 18
A. Preprocessing of the *E. coli* and mammalian data 18
B. Fitting the experimental data 18
C. Bootstrap simulations 20
D. Refitting the simulations and confidence intervals 20

XIII. Additional *E. coli* datasets 20

References 24

*
†
‡

#### I. MODEL AND DEFINITIONS

We consider a single cell cycle that begins at birth, $t = 0$, with cell volume $v_b$, and ends at division, $t = \tau$, with division volume $v_d$. The instantaneous growth rate $\lambda(t)$ determines the cell volume through

$$v(t) = v_b \exp\left(\int_0^t \lambda(t')\, dt'\right). \tag{1}$$

It is convenient to define the accumulated growth since birth,

$$I(t) \equiv \int_0^t \lambda(t')\, dt', \tag{2}$$

so that

$$v(t) = v_b e^{I(t)}. \tag{3}$$

To model cell size regulation, we use the size regulation model introduced in Ref. [1]. The division size is set by

$$v_d = 2\Delta\alpha + 2\,(1-\alpha)\, v_b + \xi, \tag{4}$$

where $\Delta$ is the mean birth size, $0 < \alpha < 2$ controls the size regulation strategy, and $\xi$ is a Gaussian noise term with zero mean and variance $\sigma_\xi^2$. The division time is determined implicitly by the growth trajectory through

$$\log \frac{v_d}{v_b} = I(\tau) = \int_0^\tau \lambda(t')\, dt'. \tag{5}$$

During a cell cycle, the biological growth rate follows an Ornstein-Uhlenbeck (OU) process,

$$\dot{\lambda}(t) = -\gamma\left(\lambda(t) - \lambda_0^{(L)}\right) + \sqrt{2D_\lambda}\,\eta(t), \quad \langle \eta(t)\rangle = 0, \quad \langle \eta(t)\eta(t')\rangle = \delta(t-t'). \tag{6}$$

Here $\gamma = 1/\tau_{\mathrm{rel}}$ is the relaxation rate, $D_\lambda$ sets the strength of the continuous fluctuations, and $\lambda_0^{(L)}$ is the mean growth rate of lineage $L$.

To account for lineage-to-lineage variability, we assume that each lineage $L$ has its own mean growth rate,

$$\lambda_0^{(L)} = \lambda_0 + \ell_L, \qquad \ell_L \sim \mathcal{N}(0, D_\ell), \tag{7}$$

where $\lambda_0$ is the population mean and $D_\ell$ is the variance across lineages.

To allow for perturbations at cell division, we also include division kicks. For the cycle $j+1$ which is born from cycle $j$, we write

$$\lambda^{(j+1)}(0) = \lambda^{(j)}(\tau_j) + \eta_d^{(j)}, \qquad \eta_d^{(j)} \sim \mathcal{N}(0, D_d), \tag{8}$$

so that $D_d$ is the variance of the division-associated perturbations, which we term “division kicks”.

For measurements taken at discrete times $t_k = k\Delta t$, the true growth rate is

$$\lambda_k \equiv \frac{1}{\Delta t} \log \frac{v_{k+1}}{v_k} = \frac{1}{\Delta t} \int_{t_k}^{t_{k+1}} \lambda(t')\, dt', \tag{9}$$

and the accumulated growth up to time $t_k$ is

$$I_k \equiv I(t_k) = \log \frac{v_k}{v_b}. \tag{10}$$

The instantaneous growth rate $\lambda_k$ and the accumulated growth rate $I_k$ are the biological observables whose autocovariances we derive below.

#### II. CONTINUOUS-TIME AUTOCOVARIANCE OF THE BIOLOGICAL GROWTH RATE

We now derive the autocovariance of $\lambda(t)$ within a cell cycle, including the effects of continuous OU fluctuations, division kicks, and lineage-to-lineage variability. In this section, all quantities correspond to continuous time, where the process $\lambda(t)$ is defined. In the next section, we consider the values taken at discrete times which correspond to measured quantities.

For a fixed lineage $L$, with mean growth rate $\lambda_0 + \ell_L$, the OU equation is

$$\dot{\lambda}(t) = -\gamma\left(\lambda(t) - \lambda_0 - \ell_L\right) + \sqrt{2D_\lambda}\,\eta(t). \tag{11}$$

Given the growth rate at birth $\lambda_b \equiv \lambda(t=0)$, its solution is

$$\lambda(t) - \lambda_0 - \ell_L = \left(\lambda_b - \lambda_0 - \ell_L\right)e^{-\gamma t} + \sqrt{2D_\lambda}\int_0^t e^{-\gamma(t-t')}\eta(t')\,dt'. \tag{12}$$

We define the integrated noise term as

$$\zeta(t) \equiv \int_0^t e^{-\gamma(t-t')}\eta(t')\,dt', \qquad \langle\zeta(t)\rangle = 0, \qquad \mathrm{Var}\left(\zeta(t)\right) = \frac{1-e^{-2\gamma t}}{2\gamma}. \tag{13}$$

and the solution can be written as

$$\lambda(t) - \lambda_0 - \ell_L = \left(\lambda_b - \lambda_0 - \ell_L\right)e^{-\gamma t} + \sqrt{2D_\lambda}\,\zeta(t). \tag{14}$$

From this, we can compute the autocovariance of $\lambda(t)$ at two different times $t'$ and $t''$,

$$\mathrm{Cov}\left(\lambda(t'), \lambda(t'')|\ell_L\right) = \frac{D_\lambda}{\gamma}e^{-\gamma|t''-t'|} + \left(\mathrm{Var}\left[\lambda_b|\ell_L\right] - \frac{D_\lambda}{\gamma}\right)e^{-\gamma(t'+t'')}. \tag{15}$$

This expression depends explicitly on the variance of the birth growth rate $\lambda_b$ across cells from the same lineage, which we compute next.

The statistics of the birth growth rate are determined by the division kicks and the OU relaxation during the previous cell cycle. Across generations, going from cycle $j$ to cycle $j+1$, the birth growth rate obeys

$$\lambda_b^{(j+1)} - \lambda_0 - \ell_L = \left(\lambda_b^{(j)} - \lambda_0 - \ell_L\right)e^{-\gamma\tau_j} + z_j, \qquad z_j = \sqrt{2D_\lambda}\,\zeta(\tau_j) + \eta_d^{(j)}, \tag{16}$$

and the variance of the combined noise $z_j$ is

$$\mathrm{Var}\left(z_j|\tau_j\right) = \frac{D_\lambda}{\gamma}\left(1 - e^{-2\gamma\tau_j}\right) + D_d. \tag{17}$$

To obtain a closed form expression for $\mathrm{Var}\left(\lambda_b|\ell_L\right)$, we approximate the random division time by its mean value $\langle\tau\rangle$. Then the birth process becomes an autoregressive process with coefficient $e^{-\gamma\langle\tau\rangle}$,

$$\lambda_b^{(j+1)} - \lambda_0 - \ell_L = e^{-\gamma\langle\tau\rangle}\left(\lambda_b^{(j)} - \lambda_0 - \ell_L\right) + z_j, \tag{18}$$

and its stationary variance is

$$\mathrm{Var}\left(\lambda_b|\ell_L\right) \approx \frac{D_\lambda}{\gamma} + \frac{D_d}{1 - e^{-2\gamma\langle\tau\rangle}}. \tag{19}$$

Importantly, this conditional variance does not depend on $\ell_L$.

Substituting this result into Eq. (15) results in

$$\mathrm{Cov}\left(\lambda(t'), \lambda(t'')|\ell_L\right) = \frac{D_\lambda}{\gamma}e^{-\gamma|t''-t'|} + \frac{D_d}{1 - e^{-2\gamma\langle\tau\rangle}}e^{-\gamma(t'+t'')}. \tag{20}$$

We now average over lineages. Using the law of total autocovariance,

$$\mathrm{Cov}\left(\lambda(t'),\lambda(t'')\right)=\mathbb{E}_\ell\left[\mathrm{Cov}\left(\lambda(t'),\lambda(t'')|\ell_L\right)\right]+\mathrm{Cov}_\ell\left(\mathbb{E}[\lambda(t')|\ell_L],\mathbb{E}[\lambda(t'')|\ell_L]\right). \tag{21}$$

For a given lineage, the mean growth rate is $\lambda_0+\ell_L$, so

$$\mathbb{E}[\lambda(t)|\ell_L]=\lambda_0+\ell_L. \tag{22}$$

Therefore the second term in Eq. (21) is simply

$$\mathrm{Cov}_\ell\left(\lambda_0+\ell_L,\lambda_0+\ell_L\right)=D_\ell. \tag{23}$$

We thus obtain the continuous-time autocovariance of the biological growth rate (substituting Eqs. (20), (23) into Eq. (21)),

$$\mathrm{Cov}\left(\lambda(t'),\lambda(t'')\right)=\frac{D_\lambda}{\gamma}e^{-\gamma|t''-t'|}+\frac{D_d}{1-e^{-2\gamma\langle\tau\rangle}}e^{-\gamma(t'+t'')}+D_\ell. \tag{24}$$

Note that in the main text we have considered the case $\gamma\langle\tau\rangle\gg 1$ and therefore the division kicks contribution was approximated as $D_d/\left(1-e^{-2\gamma\langle\tau\rangle}\right)\approx D_d$.

This expression shows that the three biological mechanisms contribute in qualitatively different ways. Continuous OU fluctuations generate a stationary exponentially decaying autocovariance in the time difference $|t''-t'|$. Division kicks generate a positive contribution that is strongest near birth and decays as the memory of the kick relaxes during the cell cycle. Lineage variability contributes a constant offset, reflecting the fact that cells from different lineages fluctuate around different mean growth rates.

#### III. STATISTICS OF THE INSTANTANEOUS GROWTH RATE

We now derive the autocovariance of the instantaneous growth rate $\lambda_k$, Eq. (9). Note that $\lambda_k$ is not instantaneous, but is the average of $\lambda(t)$ over the interval $(t_k,t_{k+1})$. We define the dimensionless autocovariance at lag $n$ by

$$\mathcal{A}_n(k)\equiv(\Delta t)^2\,\mathrm{Cov}\left(\lambda_k,\lambda_{k+n}\right). \tag{25}$$

Using Eq. (24), this can be written as

$$\mathcal{A}_n(k)=\int_{t_k}^{t_{k+1}}dt'\int_{t_{k+n}}^{t_{k+n+1}}dt''\,\mathrm{Cov}\left(\lambda(t'),\lambda(t'')\right)\equiv\mathcal{A}_n^{\mathrm{OU}}(k)+\mathcal{A}_n^{\mathrm{div}}(k)+\mathcal{A}_n^{\ell}(k). \tag{26}$$

The OU contribution is

$$\mathcal{A}_n^{\mathrm{OU}}(k)=\frac{D_\lambda}{\gamma}\int_{t_k}^{t_{k+1}}dt'\int_{t_{k+n}}^{t_{k+n+1}}dt''\,e^{-\gamma|t''-t'|}. \tag{27}$$

For $n=0$, this gives

$$\begin{aligned}\mathcal{A}_0^{\mathrm{OU}}(k)&=\frac{D_\lambda}{\gamma}\int_{t_k}^{t_{k+1}}dt'\int_{t_k}^{t_{k+1}}dt''\,e^{-\gamma|t''-t'|}=2\frac{D_\lambda}{\gamma}\int_{t_k}^{t_{k+1}}dt'\int_{t'}^{t_{k+1}}dt''\,e^{-\gamma(t''-t')}\\&=2\frac{D_\lambda}{\gamma^3}\left[\gamma\Delta t-\left(1-e^{-\gamma\Delta t}\right)\right].\end{aligned} \tag{28}$$

For $n\geq 1$, we have $t''\geq t'$, and therefore

$$\mathcal{A}_n^{\mathrm{OU}}(k)=\frac{D_\lambda}{\gamma}\int_{t_k}^{t_{k+1}}dt'\int_{t_{k+n}}^{t_{k+n+1}}dt''\,e^{-\gamma(t''-t')}=\frac{D_\lambda}{\gamma^3}e^{-\gamma(n-1)\Delta t}\left(1-e^{-\gamma\Delta t}\right)^2. \tag{29}$$

Thus the OU contribution is given by

$$\mathcal{A}_n^{\mathrm{OU}}(k) = \frac{D_\lambda}{\gamma^3} \begin{cases} 2\left[\gamma\Delta t - \left(1 - e^{-\gamma\Delta t}\right)\right], & n = 0, \\ e^{-\gamma(n-1)\Delta t}\left(1 - e^{-\gamma\Delta t}\right)^2, & n \geq 1. \end{cases} \tag{30}$$

The contribution of division kicks is

$$\mathcal{A}_n^{\mathrm{div}}(k) = \frac{D_d}{1 - e^{-2\gamma\langle\tau\rangle}} \int_{t_k}^{t_{k+1}} dt' \int_{t_{k+n}}^{t_{k+n+1}} dt''\, e^{-\gamma(t'+t'')} = \frac{D_d}{\gamma^2\left(1 - e^{-2\gamma\langle\tau\rangle}\right)} e^{-\gamma(2k+n)\Delta t}\left(1 - e^{-\gamma\Delta t}\right)^2. \tag{31}$$

Finally, the lineage contribution is obtained by integrating the constant term $D_\ell$,

$$\mathcal{A}_n^{\ell}(k) = D_\ell \int_{t_k}^{t_{k+1}} dt' \int_{t_{k+n}}^{t_{k+n+1}} dt'' = D_\ell(\Delta t)^2. \tag{32}$$

The three biological contributions have qualitatively different structures. The OU contribution, Eq. (30), is independent of $k$ and decays with lag $n$. The division kicks contribution, Eq. (31), decays both with age $k$ and lag $n$. The lineage contribution, Eq. (32), is a constant.

#### IV. STATISTICS OF THE ACCUMULATED GROWTH RATE

We now derive the autocovariance of the accumulated growth rate

$$I_k = \int_0^{t_k} \lambda(t')\, dt'. \tag{33}$$

We define

$$\mathcal{B}_n(k) \equiv \mathrm{Cov}\left(I_k, I_{k+n}\right). \tag{34}$$

Using Eq. (24), this can be written as

$$\mathcal{B}_n(k) = \int_0^{t_k} dt' \int_0^{t_{k+n}} dt''\, \mathrm{Cov}\left(\lambda(t'), \lambda(t'')\right) \equiv \mathcal{B}_n^{\mathrm{OU}}(k) + \mathcal{B}_n^{\mathrm{div}}(k) + \mathcal{B}_n^{\ell}(k). \tag{35}$$

The OU contribution is

$$\mathcal{B}_n^{\mathrm{OU}}(k) = \frac{D_\lambda}{\gamma} \int_0^{t_k} dt' \int_0^{t_{k+n}} dt''\, e^{-\gamma|t''-t'|}. \tag{36}$$

Since $t_{k+n} \geq t_k$, we split the integral into the regions $t'' < t'$, $t'' > t'$ with $t'' < t_k$, and $t'' > t_k$. This gives

$$\mathcal{B}_n^{\mathrm{OU}}(k) = \frac{D_\lambda}{\gamma} \int_0^{t_k} dt' \left[\int_0^{t'} dt''\, e^{-\gamma(t'-t'')} + \int_{t'}^{t_k} dt''\, e^{-\gamma(t''-t')} + \int_{t_k}^{t_{k+n}} dt''\, e^{-\gamma(t''-t')}\right]. \tag{37}$$

Evaluating the three terms, we obtain

$$\mathcal{B}_n^{\mathrm{OU}}(k) = 2\frac{D_\lambda}{\gamma^3}\left(\gamma t_k - 1 + e^{-\gamma t_k}\right) + \frac{D_\lambda}{\gamma^3}\left(1 - e^{-\gamma t_k}\right)\left(1 - e^{-\gamma(t_{k+n} - t_k)}\right). \tag{38}$$

The contribution of division kicks is

$$\mathcal{B}_n^{\mathrm{div}}(k) = \frac{D_d}{1 - e^{-2\gamma\langle\tau\rangle}} \int_0^{t_k} dt' \int_0^{t_{k+n}} dt''\, e^{-\gamma(t'+t'')} = \frac{D_d}{\gamma^2\left(1 - e^{-2\gamma\langle\tau\rangle}\right)}\left(1 - e^{-\gamma t_k}\right)\left(1 - e^{-\gamma t_{k+n}}\right). \tag{39}$$

Finally, the lineage contribution is obtained by integrating the constant term $D_\ell$,

$$\mathcal{B}_n^\ell(k) = D_\ell \int_0^{t_k} dt' \int_0^{t_{k+n}} dt'' = D_\ell\, t_k t_{k+n}. \tag{40}$$

As for $\mathcal{A}_n(k)$ in the previous section, the three biological contributions have distinct time dependence. At late times $\gamma t \gg 1$, the OU contribution (38) grows linearly with age, the division kick contribution (39) approaches a constant plateau, and the lineage contribution (40) grows quadratically with age. This difference is what makes $\mathcal{B}_n(k)$, and in particular $\mathcal{B}_1(k)$, useful for distinguishing between different biological sources of growth rate variability.

#### V. MEASUREMENT NOISE

We now derive the contribution of measurement noise to the instantaneous growth rate and accumulated growth autocovariances.

We consider additive, Gaussian, and uncorrelated measurement noise in the cell volume,

$$\hat{v}_k = v_k + \epsilon_k, \qquad \epsilon_k \sim \mathcal{N}(0, D_\epsilon), \qquad \langle \epsilon_k \epsilon_{k'} \rangle = D_\epsilon \delta_{k,k'}. \tag{41}$$

We assume that the measurement noise is independent of the biological dynamics and small compared with the cell volume, $\epsilon_k / v_k \ll 1$.

The measured growth rate is (recall Eq. (9))

$$\hat{\lambda}_k \equiv \frac{1}{\Delta t} \log \frac{\hat{v}_{k+1}}{\hat{v}_k}. \tag{42}$$

Expanding the logarithm to leading order in $\epsilon_k / v_k$, we have

$$\begin{aligned} \hat{\lambda}_k &= \frac{1}{\Delta t} \log \frac{v_{k+1} + \epsilon_{k+1}}{v_k + \epsilon_k} = \frac{1}{\Delta t} \left[ \log \frac{v_{k+1}}{v_k} + \log \left( \frac{1 + \frac{\epsilon_{k+1}}{v_{k+1}}}{1 + \frac{\epsilon_k}{v_k}} \right) \right] \\ &= \lambda_k + \frac{1}{\Delta t} \left[ \log \left( 1 + \frac{\epsilon_{k+1}}{v_{k+1}} \right) - \log \left( 1 + \frac{\epsilon_k}{v_k} \right) \right] = \lambda_k + \frac{1}{\Delta t} \left[ \frac{\epsilon_{k+1}}{v_{k+1}} - \frac{\epsilon_k}{v_k} + \mathcal{O}\left( \left( \frac{\epsilon}{v} \right)^2 \right) \right]. \end{aligned} \tag{43}$$

Therefore

$$\hat{\lambda}_k = \lambda_k + \delta\hat{\lambda}_k + \mathcal{O}\left[ \left( \frac{\epsilon}{v} \right)^2 \right], \tag{44}$$

where

$$\delta\hat{\lambda}_k = \frac{1}{\Delta t} \left( \frac{\epsilon_{k+1}}{v_{k+1}} - \frac{\epsilon_k}{v_k} \right). \tag{45}$$

Since the measurement noise is independent of the biological dynamics and has zero mean, we have

$$\mathbb{E}\left[ \delta\hat{\lambda}_k \right] = 0. \tag{46}$$

The autocovariance of the measurement noise contribution is therefore

$$\mathrm{Cov}\left( \delta\hat{\lambda}_k, \delta\hat{\lambda}_{k+n} \right) = \mathbb{E}\left[ \delta\hat{\lambda}_k\, \delta\hat{\lambda}_{k+n} \right]. \tag{47}$$

Substituting Eq. (45), into Eq. (47) we find

$$\mathbb{E}\left[ \delta\hat{\lambda}_k\, \delta\hat{\lambda}_{k+n} \right] = \frac{1}{(\Delta t)^2} \mathbb{E}\left[ \left( \frac{\epsilon_{k+1}}{v_{k+1}} - \frac{\epsilon_k}{v_k} \right) \left( \frac{\epsilon_{k+n+1}}{v_{k+n+1}} - \frac{\epsilon_{k+n}}{v_{k+n}} \right) \right]. \tag{48}$$

Because the measurement noise is independent at different times, only terms with matching indices survive. As a result,

$$\mathrm{Cov}\left(\delta\hat{\lambda}_k, \delta\hat{\lambda}_{k+n}\right) = \frac{D_\epsilon}{(\Delta t)^2}\begin{cases}\left\langle v_k^{-2}\right\rangle + \left\langle v_{k+1}^{-2}\right\rangle, & n=0,\\ -\left\langle v_{k+1}^{-2}\right\rangle, & n=1,\\ 0, & n\geq 2.\end{cases} \tag{49}$$

Therefore, we find that the corresponding contribution to the instantaneous growth rate autocovariance is

$$\delta\hat{\mathcal{A}}_n(k) \equiv (\Delta t)^2\,\mathrm{Cov}\left(\delta\hat{\lambda}_k, \delta\hat{\lambda}_{k+n}\right) = D_\epsilon \times \begin{cases}\left\langle v_k^{-2}\right\rangle + \left\langle v_{k+1}^{-2}\right\rangle, & n=0,\\ (-1)\left\langle v_{k+1}^{-2}\right\rangle, & n=1,\\ 0, & n\geq 2.\end{cases} \tag{50}$$

We next consider the measured accumulated growth

$$\hat{I}_k \equiv \log\frac{\hat{v}_k}{\hat{v}_b} = I_k + \delta\hat{I}_k + \mathcal{O}\left[\left(\frac{\epsilon}{v}\right)^2\right], \tag{51}$$

where

$$\delta\hat{I}_k = \frac{\epsilon_k}{v_k} - \frac{\epsilon_b}{v_b}. \tag{52}$$

Again, $\mathbb{E}\left[\delta\hat{I}_k\right] = 0$, and the autocovariance is

$$\mathrm{Cov}\left(\delta\hat{I}_k, \delta\hat{I}_{k+n}\right) = \mathbb{E}\left[\delta\hat{I}_k\,\delta\hat{I}_{k+n}\right]. \tag{53}$$

Substituting Eq. (52) into Eq. (53), we obtain

$$\mathbb{E}\left[\delta\hat{I}_k\,\delta\hat{I}_{k+n}\right] = \mathbb{E}\left[\left(\frac{\epsilon_k}{v_k} - \frac{\epsilon_b}{v_b}\right)\left(\frac{\epsilon_{k+n}}{v_{k+n}} - \frac{\epsilon_b}{v_b}\right)\right]. \tag{54}$$

For $n \geq 1$, the only surviving contribution comes from the common birth term $\epsilon_b$, whereas for $n = 0$ there is an additional contribution from $\epsilon_k$. Thus,

$$\delta\hat{\mathcal{B}}_n(k) \equiv \mathrm{Cov}\left(\delta\hat{I}_k, \delta\hat{I}_{k+n}\right) = D_\epsilon\left(\left\langle v_b^{-2}\right\rangle + \delta_{n,0}\left\langle v_k^{-2}\right\rangle\right). \tag{55}$$

This implies that for every lag $n \geq 1$, the measurement noise contribution to $\hat{\mathcal{B}}_n(k)$ is simply the constant

$$\delta\hat{\mathcal{B}}_{n\geq 1}(k) = D_\epsilon\left\langle v_b^{-2}\right\rangle. \tag{56}$$

To obtain a simple approximation for $\left\langle v_k^{-2}\right\rangle$, we use

$$v(t) = v_b e^{I(t)}. \tag{57}$$

Since $\lambda(t)$ is a Gaussian process, $I(t) = \int_0^t \lambda(t')dt'$ is also Gaussian, with mean $\langle I(t)\rangle = \lambda_0 t$ and variance $\mathrm{Var}\,(I(t))$. Therefore,

$$\left\langle e^{-2I(t)}\right\rangle = e^{-2\lambda_0 t + 2\,\mathrm{Var}(I(t))}, \tag{58}$$

and thus

$$\left\langle v^{-2}(t)\right\rangle = \left\langle v_b^{-2} e^{-2I(t)}\right\rangle. \tag{59}$$

Within our model, $v_b$ and the subsequent growth trajectory $\lambda(t)$ (and hence $I(t) = \int_0^t \lambda(t')dt'$) are uncorrelated and therefore

$$\left\langle v^{-2}(t)\right\rangle = \left\langle v_b^{-2}\right\rangle e^{-2\lambda_0 t + 2\,\mathrm{Var}(I(t))}. \tag{60}$$

If the variance correction $\mathrm{Var}(I(t))$ is small with respect to $\lambda_0 t$, this further reduces to

$$\left\langle v_k^{-2}\right\rangle \approx \left\langle v_b^{-2}\right\rangle e^{-2\lambda_0 k\Delta t}. \tag{61}$$

Substituting this approximation into Eq. (50), and assuming $\lambda_0 \Delta t \ll 1$, gives the simple relation

$$\delta\hat{\mathcal{A}}_n(k) \approx D_\epsilon \left\langle v_b^{-2}\right\rangle e^{-2\lambda_0 k\Delta t} \times \begin{cases} 2, & n = 0,\\ (-1), & n = 1,\\ 0, & n \geq 2. \end{cases} \tag{62}$$

Thus, the measurement noise contribution to $\hat{\mathcal{A}}_n(k)$ has a characteristic lag structure: it is positive at lag 0, negative at lag 1, and vanishes for lags $n \geq 2$. By contrast, its contribution to $\hat{\mathcal{B}}_n(k)$, Eq. (55), is simpler as for $n \geq 1$ it is independent of both $k$ and $n$.

##### A. Alternative centered finite-difference estimator

In the main text we define the inferred growth rate using a forward finite difference, as in Eq. (42). A natural alternative is the centered finite-difference estimator (denoted with the superscript $(c)$)

$$\hat{\lambda}_k^{(c)} \equiv \frac{1}{2\Delta t} \log \frac{\hat{v}_{k+1}}{\hat{v}_{k-1}}. \tag{63}$$

Expanding the logarithm to leading order in $\epsilon_k / v_k$, we obtain

$$\hat{\lambda}_k^{(c)} = \frac{1}{2\Delta t} \log \frac{v_{k+1}}{v_{k-1}} + \delta\hat{\lambda}_k^{(c)} + \mathcal{O}\left[\left(\frac{\epsilon}{v}\right)^2\right], \qquad \delta\hat{\lambda}_k^{(c)} = \frac{1}{2\Delta t}\left(\frac{\epsilon_{k+1}}{v_{k+1}} - \frac{\epsilon_{k-1}}{v_{k-1}}\right). \tag{64}$$

Since the measurement noise has zero mean and is independent of the biological dynamics, the corresponding autocovariance is

$$\mathrm{Cov}\left(\delta\hat{\lambda}_k^{(c)}, \delta\hat{\lambda}_{k+n}^{(c)}\right) = \mathbb{E}\left[\delta\hat{\lambda}_k^{(c)} \delta\hat{\lambda}_{k+n}^{(c)}\right].$$

Substituting Eq. (64) and using the independence of the measurement noise at different times, we find

$$\mathrm{Cov}\left(\delta\hat{\lambda}_k^{(c)}, \delta\hat{\lambda}_{k+n}^{(c)}\right) = \frac{D_\epsilon}{4(\Delta t)^2} \begin{cases} \left\langle v_{k-1}^{-2}\right\rangle + \left\langle v_{k+1}^{-2}\right\rangle, & n = 0,\\ 0, & n = 1,\\ -\left\langle v_{k+1}^{-2}\right\rangle, & n = 2,\\ 0, & n \geq 3. \end{cases} \tag{65}$$

Therefore, the contribution of measurement noise to the instantaneous growth rate autocovariance becomes

$$\delta\hat{\mathcal{A}}_n^{(c)}(k) \equiv (\Delta t)^2\, \mathrm{Cov}\left(\delta\hat{\lambda}_k^{(c)}, \delta\hat{\lambda}_{k+n}^{(c)}\right) = \frac{D_\epsilon}{4} \begin{cases} \left\langle v_{k-1}^{-2}\right\rangle + \left\langle v_{k+1}^{-2}\right\rangle, & n = 0,\\ 0, & n = 1,\\ -\left\langle v_{k+1}^{-2}\right\rangle, & n = 2,\\ 0, & n \geq 3. \end{cases} \tag{66}$$

Using the approximation in Eq. (61), and assuming $\lambda_0 \Delta t \ll 1$, this reduces to

$$\delta\hat{\mathcal{A}}_n^{(c)}(k) \approx D_\epsilon \left\langle v_b^{-2} \right\rangle e^{-2\lambda_0 k \Delta t} \begin{cases} \frac{1}{2}, & n = 0, \\ 0, & n = 1, \\ -\frac{1}{4}, & n = 2, \\ 0, & n \geq 3. \end{cases} \tag{67}$$

Thus, for the centered estimator the measurement noise contribution to $\hat{\mathcal{A}}_n(k)$ remains qualitatively similar to the forward estimator, Eq. (62), but its lag structure is shifted: it is positive at lag 0, vanishes at lag 1, becomes negative at lag 2, and vanishes for $n \geq 3$. By contrast, the measurement noise contribution to $\hat{\mathcal{B}}_n(k)$ is unchanged, since $\hat{I}_k = \log(\hat{v}_k/\hat{v}_b)$ does not depend on the finite-difference convention used to define the inferred instantaneous growth rate.

#### VI. ASYMPTOTIC BEHAVIOR OF $\mathcal{B}_1(k)$

We now derive the early-time and late-time behavior of $\mathcal{B}_1(k)$, which is used in the main text to distinguish between different biological sources of growth rate variability.

For $n = 1$, Eqs. (38), (39), and (40) give

$$\mathcal{B}_1^{\mathrm{OU}}(k) = 2\frac{D_\lambda}{\gamma^3}\left(\gamma t_k - 1 + e^{-\gamma t_k}\right) + \frac{D_\lambda}{\gamma^3}\left(1 - e^{-\gamma t_k}\right)\left(1 - e^{-\gamma \Delta t}\right), \tag{68a}$$

$$\mathcal{B}_1^{\mathrm{div}}(k) = \frac{D_d}{\gamma^2\left(1 - e^{-2\gamma\langle\tau\rangle}\right)}\left(1 - e^{-\gamma t_k}\right)\left(1 - e^{-\gamma(t_k + \Delta t)}\right), \tag{68b}$$

$$\mathcal{B}_1^{\ell}(k) = D_\ell\, t_k\,(t_k + \Delta t)\,. \tag{68c}$$

We first consider early times, $t_k \ll 1/\gamma$, and assume $\gamma \Delta t \ll 1$. Expanding to leading order, we obtain

$$\mathcal{B}_1^{\mathrm{OU}}(k) \approx \frac{D_\lambda}{\gamma} t_k\,(t_k + \Delta t) \approx \frac{D_\lambda}{\gamma} t_k^2, \tag{69a}$$

$$\mathcal{B}_1^{\mathrm{div}}(k) \approx \frac{D_d}{1 - e^{-2\gamma\langle\tau\rangle}} t_k\,(t_k + \Delta t) \approx \frac{D_d}{1 - e^{-2\gamma\langle\tau\rangle}} t_k^2, \tag{69b}$$

$$\mathcal{B}_1^{\ell}(k) = D_\ell\, t_k\,(t_k + \Delta t) \approx D_\ell\, t_k^2. \tag{69c}$$

Thus, at early times all three contributions have the same quadratic dependence on age. They differ only by their prefactors. This implies that the early-time behavior of $\mathcal{B}_1(k)$ alone is not sufficient to distinguish between these biological mechanisms.

We now consider late times, $t_k \gg 1/\gamma$. In this regime, the exponentials $e^{-\gamma t_k}$ and $e^{-\gamma(t_k + \Delta t)}$ are negligible. Since $\langle\tau\rangle \gtrsim t_k$, we may also neglect $e^{-2\gamma\langle\tau\rangle}$, and we obtain the leading behavior

$$\mathcal{B}_1^{\mathrm{OU}}(k) \approx 2\frac{D_\lambda}{\gamma^2} t_k, \tag{70a}$$

$$\mathcal{B}_1^{\mathrm{div}}(k) \approx \frac{D_d}{\gamma^2}, \tag{70b}$$

$$\mathcal{B}_1^{\ell}(k) \approx D_\ell\, t_k^2. \tag{70c}$$

Therefore, at late times the three mechanisms become distinguishable by their time trends. Continuous OU fluctuations generate a contribution that grows linearly with age, division kicks generate a constant plateau, and finally, lineage variability generates a contribution that grows quadratically with age. This separation of asymptotic behavior is what makes $\mathcal{B}_1(k)$ a useful observable for distinguishing between different sources of biological growth rate variability.

#### VII. SURVIVAL BIAS IN AGE-CONDITIONED STATISTICS

The statistics analyzed in the main text, $\hat{\mathcal{A}}_n(k), \hat{\mathcal{B}}_n(k)$, are functions of the cell age since birth. At a given observation time $T$, the statistics are therefore computed only over cells that have not yet divided by age $T$, which is the origin of survival bias.

To see how to correct for it, consider the finite-sample case, where we have a total of $N$ cell cycles, and each cycle has a division time $\tau_i$. Let $O_i(T)$ be some observable defined at time $T$ for the $i$-th cell cycle. Then, the empirical average at age $T$ is obtained by averaging only over the cells that have not yet divided by age $T$,

$$\langle O(T)\rangle_N = \frac{\sum_{i=1}^N O_i(T)\mathbf{1}_{\tau_i>T}}{\sum_{i=1}^N \mathbf{1}_{\tau_i>T}} = \frac{1}{N_T}\sum_{i=1}^N O_i(T)\,\mathbf{1}_{\{\tau_i>T\}}, \tag{71}$$

where $\mathbf{1}_{\tau_i>T}$ is equal to 1 if the $i$-th cell has not divided up to time $T$, and 0 otherwise. The denominator $N_T = \sum_{i=1}^N \mathbf{1}_{\{\tau_i>T\}}$ is the number of cells that have not yet divided by age $T$. In the infinite sample limit, this empirical average converges to the conditional expectation

$$\mathbb{E}\left[O|\tau > T\right] = \frac{\mathbb{E}\left[O\,\mathbf{1}_{\{\tau>T\}}\right]}{P\left(\tau > T\right)}. \tag{72}$$

This is the quantity that we need to compute in order to correct for survival bias.

Let us now write the survival event $\{\tau > T\}$ in terms of the accumulated growth $I(T)$. Using $v(T) = v_b e^{I(T)}$, the survival event can be written as

$$\{\tau > T\} = \{v(T) < v_d\} = \{\log v(T) < \log v_d\} = \{I(T) < L_{\text{th}}\}, \tag{73}$$

where we denote the log-threshold $L_{\text{th}}$, using Eq. (4),

$$L_{\text{th}} \equiv \log\frac{v_d}{v_b} = \log\frac{2\alpha\Delta + 2\left(1-\alpha\right)v_b + \xi}{v_b}. \tag{74}$$

The denominator in Eq. (72) is the survival probability

$$P\left(\tau > T\right) = P\left(I(T) < L_{\text{th}}\right). \tag{75}$$

To evaluate it, we first fix $v_b$ and $\xi$, so that $L_{\text{th}}$ is fixed. As the random variable $I(T)$ is Gaussian, with mean $\mu_I(T)$ and variance $\sigma_I^2(T)$, we have

$$P\left(\tau > T|v_b,\xi\right) = P\left(I(T) < L_{\text{th}}|v_b,\xi\right) = \Phi\left(\frac{L_{\text{th}} - \mu_I(T)}{\sigma_I(T)}\right) = \Phi(z(T;v_b,\xi)), \tag{76}$$

$$z(T;v_b,\xi) \equiv \frac{L_{\text{th}} - \mu_I(T)}{\sigma_I(T)}, \qquad \Phi(z) = \int_{-\infty}^{z}\phi(y)dy, \qquad \phi(z) = \frac{1}{\sqrt{2\pi}}e^{-z^2/2}, \tag{77}$$

where $\Phi(z)$ is the standard normal cumulative distribution function [2]. Finally, to obtain the unconditional survival probability, we need to average over $v_b$ and $\xi$,

$$P\left(\tau > T\right) = \mathbb{E}_{v_b,\xi}\left[\Phi(z(T;v_b,\xi))\right]. \tag{78}$$

We now turn to the numerator in Eq. (72), which is the joint expectation of an observable $O$ and the survival event. Recall that the statistics $\hat{\mathcal{A}}, \hat{\mathcal{B}}$ are covariances of the instantaneous growth rate $\lambda_k$ and the accumulated growth $I_k$ (Eqs. (25), (34)). We therefore consider two generic observables $X$ and $Y$ that are jointly Gaussian with $I(T)$, and derive the survival conditioned covariance in this general setting.

We first derive the result for a fixed threshold $L_{\text{th}} = L$, and then average over $v_b$ and $\xi$ at the end. At fixed $L$, the event that the cell has not yet divided by age $T$ is

$$A_{T,L} \equiv \{I(T) < L\}. \tag{79}$$

For $X, Y$, jointly Gaussian with $I(T)$, we want to compute

$$\text{Cov}\left(X(T), Y(T)|A_{T,L}\right) = \text{Cov}\left(X(T), Y(T)|I(T) < L\right). \tag{80}$$

Let us denote

$$\mu_X = \mathbb{E}\left[X(T)\right], \qquad \mu_Y = \mathbb{E}\left[Y(T)\right], \qquad \mu_I = \mathbb{E}\left[I(T)\right], \tag{81}$$

and

$$c_X = \mathrm{Cov}\left(X(T), I(T)\right), \qquad c_Y = \mathrm{Cov}\left(Y(T), I(T)\right), \qquad \sigma_I^2 = \mathrm{Var}\left(I(T)\right). \tag{82}$$

Since $X(T)$, $Y(T)$, and $I(T)$ are jointly Gaussian, we can decompose

$$X(T) = \mu_X + a_X\left(I(T) - \mu_I\right) + \varepsilon_X, \qquad a_X \equiv \frac{c_X}{\sigma_I^2} \tag{83a}$$

$$Y(T) = \mu_Y + a_Y\left(I(T) - \mu_I\right) + \varepsilon_Y, \qquad a_Y \equiv \frac{c_Y}{\sigma_I^2} \tag{83b}$$

where the residuals $\varepsilon_X$ and $\varepsilon_Y$ are Gaussian and independent of $I(T)$, as, by construction,

$$\mathrm{Cov}\left(\varepsilon_X, I(T)\right) = 0, \qquad \mathrm{Cov}\left(\varepsilon_Y, I(T)\right) = 0. \tag{84}$$

The important point is that since the survival event $A_{T,L} = \{I(T) < L\}$ depends only on $I(T)$, the residuals $\varepsilon_X$ and $\varepsilon_Y$ are also independent of the survival event $A_{T,L}$.

Now we compute the conditional covariance, (80),

$$\mathrm{Cov}\left(X(T), Y(T)|A_{T,L}\right) = \mathbb{E}\left[\left(X(T) - \mathbb{E}\left[X(T)|A_{T,L}\right]\right)\left(Y(T) - \mathbb{E}\left[Y(T)|A_{T,L}\right]\right)|A_{T,L}\right]. \tag{85}$$

We start with the conditional means $\mathbb{E}\left[X(T)|A_{T,L}\right]$ and $\mathbb{E}\left[Y(T)|A_{T,L}\right]$. From Eq. (83a), we have

$$\mathbb{E}\left[X(T)|A_{T,L}\right] = \mu_X + a_X\left(\mathbb{E}\left[I(T)|A_{T,L}\right] - \mu_I\right) + \cancel{\mathbb{E}\left[\varepsilon_X|A_{T,L}\right]}, \tag{86}$$

where the last term is zero since $\varepsilon_X$ is independent of $A_{T,L}$ and has zero mean. The same formula holds for $\mathbb{E}\left[Y(T)|A_{T,L}\right]$. Subtracting the conditional means from $X(T)$ and $Y(T)$, we find

$$X(T) - \mathbb{E}\left[X(T)|A_{T,L}\right] = a_X\left(I(T) - \mathbb{E}\left[I(T)|A_{T,L}\right]\right) + \varepsilon_X, \tag{87}$$

$$Y(T) - \mathbb{E}\left[Y(T)|A_{T,L}\right] = a_Y\left(I(T) - \mathbb{E}\left[I(T)|A_{T,L}\right]\right) + \varepsilon_Y. \tag{88}$$

Substituting into Eq. (85), we obtain

$$\mathrm{Cov}\left(X(T), Y(T)|A_{T,L}\right) = a_X a_Y \mathrm{Var}\left(I(T)|A_{T,L}\right) + \mathrm{Cov}\left(\varepsilon_X, \varepsilon_Y\right). \tag{89}$$

Indeed, the two cross terms vanish because $\varepsilon_X$ and $\varepsilon_Y$ are independent of $A_{T,L}$, and therefore also independent of $\left(I(T) - \mathbb{E}\left[I(T)|A_{T,L}\right]\right)$.

We next compute the second term $\mathrm{Cov}\left(\varepsilon_X, \varepsilon_Y\right)$. Let us express it using the original variables $X(T)$, $Y(T)$, and $I(T)$. Using Eqs. (83), we have

$$\mathrm{Cov}\left(X(T), Y(T)\right) = a_X a_Y \sigma_I^2 + \mathrm{Cov}\left(\varepsilon_X, \varepsilon_Y\right), \tag{90}$$

as the cross terms $\mathrm{Cov}\left(\varepsilon_X, I(T)\right) = \mathrm{Cov}\left(\varepsilon_Y, I(T)\right) = 0$ vanish by independence. Substituting Eq. (90) into Eq. (89), we obtain

$$\begin{aligned}\mathrm{Cov}\left(X(T), Y(T)|A_{T,L}\right) &= a_X a_Y \mathrm{Var}\left(I(T)|A_{T,L}\right) + \mathrm{Cov}\left(X(T), Y(T)\right) - a_X a_Y \sigma_I^2 \\ &= \mathrm{Cov}\left(X(T), Y(T)\right) - a_X a_Y\left[\sigma_I^2 - \mathrm{Var}\left(I(T)|A_{T,L}\right)\right]. \end{aligned} \tag{91}$$

Using the definitions of $a_X$ and $a_Y$, Eqs. (83), we can write this as

$$\mathrm{Cov}\left(X(T), Y(T)|A_{T,L}\right) = \mathrm{Cov}\left(X(T), Y(T)\right) - \frac{c_X c_Y}{\sigma_I^4}\left[\sigma_I^2 - \mathrm{Var}\left(I(T)|A_{T,L}\right)\right]. \tag{92}$$

The crucial point is that the condition on survival appears only in the term $\mathrm{Var}\left(I(T)|A_{T,L}\right)$.

It remains to evaluate the conditional variance $\mathrm{Var}\left(I(T)|A_{T,L}\right)$. For fixed $v_b$ and $\xi$, the survival probability is given in Eq. (76). We now average over the fluctuations in $v_b$ and $\xi$ that determine the threshold $L_{\rm th}$, as in Eq. (78).

We define

$$z(T; v_b, \xi) \equiv \frac{L_{\rm th} - \mu_I(T)}{\sigma_I(T)}, \tag{93}$$

and introduce

$$R_0(T) \equiv \mathbb{E}_{v_b,\xi}\left[\Phi(z)\right], \qquad R_1(T) \equiv \mathbb{E}_{v_b,\xi}\left[\phi(z)\right], \qquad R_2(T) \equiv \mathbb{E}_{v_b,\xi}\left[z\phi(z)\right], \tag{94}$$

where the expectation is taken over the distribution $P(v_b, \xi)$. The survival probability is then

$$P\left(\tau > T\right) = R_0(T). \tag{95}$$

The survival weighted first moment of $I(T)$ is

$$\mathbb{E}\left[I(T)\mathbf{1}_{\tau>T}\right] = \mathbb{E}_{v_b,\xi}\left[\mu_I(T)\Phi(z) - \sigma_I(T)\phi(z)\right] = \mu_I(T)R_0(T) - \sigma_I(T)R_1(T). \tag{96}$$

Dividing by the survival probability, Eq. (95), we obtain the conditional mean,

$$\mathbb{E}\left[I(T)|\tau > T\right] = \frac{\mathbb{E}\left[I(T)\mathbf{1}_{\tau>T}\right]}{P\left(\tau > T\right)} = \mu_I(T) - \sigma_I(T)\frac{R_1(T)}{R_0(T)}. \tag{97}$$

Similarly, the survival weighted second moment of $I(T)$ is

$$\begin{aligned}\mathbb{E}\left[I^2(T)\mathbf{1}_{\tau>T}\right] &= \mathbb{E}_{v_b,\xi}\left[\left(\mu_I^2(T) + \sigma_I^2(T)\right)\Phi(z) - 2\mu_I(T)\sigma_I(T)\phi(z) - \sigma_I^2(T)z\phi(z)\right] \\ &= \left(\mu_I^2(T) + \sigma_I^2(T)\right)R_0(T) - 2\mu_I(T)\sigma_I(T)R_1(T) - \sigma_I^2(T)R_2(T).\end{aligned} \tag{98}$$

Thus,

$$\mathbb{E}\left[I^2(T)|\tau > T\right] = \frac{\mathbb{E}\left[I^2(T)\mathbf{1}_{\tau>T}\right]}{P\left(\tau > T\right)} = \left(\mu_I^2(T) + \sigma_I^2(T)\right) - 2\mu_I(T)\sigma_I(T)\frac{R_1(T)}{R_0(T)} - \sigma_I^2(T)\frac{R_2(T)}{R_0(T)}. \tag{99}$$

Therefore, the conditional variance is

$$\begin{aligned}\mathrm{Var}\left(I(T)|\tau > T\right) &= \mathbb{E}\left[I^2(T)|A_{T,L}\right] - \left(\mathbb{E}\left[I(T)|A_{T,L}\right]\right)^2 \\ &= \sigma_I^2(T)\left[1 - \frac{R_2(T)}{R_0(T)} - \frac{R_1^2(T)}{R_0^2(T)}\right] \equiv \sigma_I^2(T)\left[1 - \chi(T)\right],\end{aligned} \tag{100}$$

Substituting this into Eq. (92), we obtain the final formula for the conditional covariance,

$$\mathrm{Cov}\left(X(T), Y(T)|A_{T,L}\right) = \mathrm{Cov}\left(X(T), Y(T)\right) - \chi(T)\frac{\mathrm{Cov}\left(X(T), I(T)\right)\mathrm{Cov}\left(Y(T), I(T)\right)}{\mathrm{Var}\left(I(T)\right)}. \tag{101}$$

##### A. Applications to $\mathcal{A}_n(k), \mathcal{B}_n(k)$

Let us apply the formula (101) to $\mathcal{B}_n(k) = \mathrm{Cov}\left(I_k, I_{k+n}\right)$. The relevant survival time is $T = t_{k+n}$. We therefore set

$$X = I_k, \qquad Y = I_{k+n}, \qquad I(T) = I_{k+n}. \tag{102}$$

Since in this case $\mathrm{Cov}\left(Y, I(T)\right) = \mathrm{Var}\left(I(T)\right)$, Eq. (101) simplifies to

$$\mathcal{B}_n^{\rm surv}(k) \equiv \mathrm{Cov}\left(I_k, I_{k+n}|\tau > t_{k+n}\right) = \left[1 - \chi(t_{k+n})\right]\mathcal{B}_n(k). \tag{103}$$

For $\mathcal{A}_n(k) = (\Delta t)^2\mathrm{Cov}\left(\lambda_k, \lambda_{k+n}\right)$, we set

$$X = \lambda_k, \qquad Y = \lambda_{k+n}, \qquad I(t_{k+n+1}) = I_{k+n+1}. \tag{104}$$

Equation (101) then gives

$$\begin{aligned}\mathcal{A}_n^{\mathrm{surv}}(k) &\equiv (\Delta t)^2 \mathrm{Cov}\left(\lambda_k, \lambda_{k+n} | \tau > t_{k+n+1}\right) \\ &= \mathcal{A}_n(k) - (\Delta t)^2 \chi(t_{k+n+1}) \frac{\mathrm{Cov}\left(\lambda_k, I_{k+n+1}\right) \mathrm{Cov}\left(\lambda_{k+n}, I_{k+n+1}\right)}{\mathrm{Var}\left(I_{k+n+1}\right)}.\end{aligned} \tag{105}$$

Equations (103) and (105) show how age-conditioned statistics differ from the unconditional expressions. The effect becomes more pronounced at late times, where the survival probability decreases and the surviving population is increasingly biased toward trajectories with smaller accumulated growth.

#### VIII. CORRELATED MEASUREMENT NOISE

In the measurement noise section above, we assumed that the additive measurement noise is independent from one time point to the next. We now generalize this to temporally correlated additive noise.

We consider additive and correlated measurement noise (compare with Eq. (41))

$$\hat{v}_k = v_k + \epsilon_k, \qquad \langle \epsilon_k \rangle = 0, \qquad \langle \epsilon_k \epsilon_{k+n} \rangle = C_n, \tag{106}$$

where $C_n$ is the autocovariance function of the measurement noise. For a stationary noise process, $C_n$ depends only on the lag $n$. As before, we assume that the measurement noise is independent of the biological dynamics.

To leading order in $\epsilon_k / v_k$, the contribution of measurement noise to the inferred growth rate is still as in (45)

$$\delta \hat{\lambda}_k = \frac{1}{\Delta t} \left( \frac{\epsilon_{k+1}}{v_{k+1}} - \frac{\epsilon_k}{v_k} \right). \tag{107}$$

Therefore,

$$\begin{aligned}\delta \hat{\mathcal{A}}_n(k) &= (\Delta t)^2 \,\mathrm{Cov}\left(\delta \hat{\lambda}_k, \delta \hat{\lambda}_{k+n}\right) \\ &= \left\langle \frac{\epsilon_{k+1}\epsilon_{k+n+1}}{v_{k+1}v_{k+n+1}} \right\rangle - \left\langle \frac{\epsilon_{k+1}\epsilon_{k+n}}{v_{k+1}v_{k+n}} \right\rangle - \left\langle \frac{\epsilon_k \epsilon_{k+n+1}}{v_k v_{k+n+1}} \right\rangle + \left\langle \frac{\epsilon_k \epsilon_{k+n}}{v_k v_{k+n}} \right\rangle \\ &= C_n \left\langle \frac{1}{v_{k+1}v_{k+n+1}} \right\rangle - C_{n-1} \left\langle \frac{1}{v_{k+1}v_{k+n}} \right\rangle - C_{n+1} \left\langle \frac{1}{v_k v_{k+n+1}} \right\rangle + C_n \left\langle \frac{1}{v_k v_{k+n}} \right\rangle.\end{aligned} \tag{108}$$

Equation (108) reduces to the result in (50) for $C_n = D_\epsilon \delta_{n,0}$.

A particularly simple approximation is obtained when the cell volume varies slowly over the correlation time of the measurement noise. Then

$$\left\langle \frac{1}{v_{k+a} v_{k+b}} \right\rangle \approx \left\langle v_k^{-2} \right\rangle, \tag{109}$$

and Eq. (108) becomes

$$\delta \hat{\mathcal{A}}_n(k) \approx \left\langle v_k^{-2} \right\rangle \left(2C_n - C_{n-1} - C_{n+1}\right). \tag{110}$$

Thus, the measurement noise contribution to $\hat{\mathcal{A}}_n(k)$ is the discrete second difference of the noise autocovariance function $C_n$. For iid noise, $C_n = \langle \epsilon_k \epsilon_{k+n} \rangle \propto \delta_{n,0}$, this gives the structure in Eq. (62): positive contribution at $n = 0$, negative contribution at $n = 1$, and zero contribution for $n \geq 2$. For correlated noise, by contrast, higher lags generally do not vanish. However, since the second difference of $C_n$ is typically largest at short lags, the measurement noise contribution to $\hat{\mathcal{A}}_n(k)$ is still expected to be most pronounced at short lags (i.e., small $n$).

We now derive the corresponding result for the accumulated growth. To leading order,

$$\delta \hat{I}_k = \frac{\epsilon_k}{v_k} - \frac{\epsilon_b}{v_b}, \tag{111}$$

and therefore

$$\delta\hat{\mathcal{B}}_n(k) \equiv \mathrm{Cov}\left(\delta\hat{I}_k, \delta\hat{I}_{k+n}\right)$$
$$= \left\langle \frac{\epsilon_k\epsilon_{k+n}}{v_k v_{k+n}} \right\rangle - \left\langle \frac{\epsilon_k\epsilon_b}{v_k v_b} \right\rangle - \left\langle \frac{\epsilon_b\epsilon_{k+n}}{v_b v_{k+n}} \right\rangle + \left\langle \frac{\epsilon_b^2}{v_b^2} \right\rangle$$
$$= C_n \left\langle \frac{1}{v_k v_{k+n}} \right\rangle - C_k \left\langle \frac{1}{v_k v_b} \right\rangle - C_{k+n} \left\langle \frac{1}{v_{k+n} v_b} \right\rangle + C_0 \left\langle v_b^{-2} \right\rangle. \tag{112}$$

Equation (112) reduces to (55) for $C_n = D_\epsilon \delta_{n,0}$. In that case, for $n \geq 1$, only the common birth term survives and $\delta\hat{\mathcal{B}}_n(k)$ is a constant. For correlated noise, this simplification is lost. Thus, temporal correlations in the measurement noise modify both observables: they generate nonzero higher-lag contributions to $\hat{\mathcal{A}}_n(k)$, and they also make the measurement noise contribution to $\hat{\mathcal{B}}_n(k)$ depend on both $k$ and $n$.

As a simple example, consider an exponentially correlated noise process,

$$C_n = D_\epsilon \rho^{|n|}, \qquad |\rho| < 1. \tag{113}$$

Using Eq. (110), we obtain

$$\delta\hat{\mathcal{A}}_0(k) \approx 2D_\epsilon (1-\rho) \left\langle v_k^{-2} \right\rangle,$$
$$\delta\hat{\mathcal{A}}_{n\geq 1}(k) \approx -D_\epsilon (1-\rho)^2 \rho^{n-1} \left\langle v_k^{-2} \right\rangle. \tag{114}$$

In this example, temporal correlations broaden the iid short-lag signature: instead of vanishing for $n \geq 2$, the measurement noise contribution extends to all lags and decays exponentially with $n$. Moreover, if the noise is positively correlated ($0 < \rho < 1$), the sign of $\delta\hat{\mathcal{A}}_n(k)$ is the same for all lags, whereas if the noise is negatively correlated ($-1 < \rho < 0$), the sign of $\delta\hat{\mathcal{A}}_n(k)$ alternates with $n$, producing an oscillatory pattern with lag.

#### IX. MULTIPLICATIVE MEASUREMENT NOISE

In some experimental settings, it may be more natural to model the measurement noise as a relative error that scales with the cell size. We therefore consider the multiplicative noise model

$$\hat{v}_k = v_k (1 + \epsilon_k), \tag{115}$$

where $\epsilon_k$ is a dimensionless random variable with

$$\langle \epsilon_k \rangle = 0, \qquad \langle \epsilon_k \epsilon_{k'} \rangle = D_\epsilon \delta_{k,k'}. \tag{116}$$

As before, we assume that the measurement noise is independent of the biological dynamics, and that $|\epsilon_k| \ll 1$.

The measured growth rate is

$$\hat{\lambda}_k \equiv \frac{1}{\Delta t} \log \frac{\hat{v}_{k+1}}{\hat{v}_k} = \frac{1}{\Delta t} \log \frac{v_{k+1}(1+\epsilon_{k+1})}{v_k (1+\epsilon_k)}. \tag{117}$$

Expanding to leading order in $\epsilon_k$, we obtain

$$\hat{\lambda}_k = \lambda_k + \delta\hat{\lambda}_k + \mathcal{O}\left(\epsilon^2\right), \qquad \delta\hat{\lambda}_k = \frac{1}{\Delta t}(\epsilon_{k+1} - \epsilon_k). \tag{118}$$

Therefore, the measurement noise contribution is given by

$$\delta\hat{\mathcal{A}}_n(k) \equiv (\Delta t)^2 \mathrm{Cov}\left(\delta\hat{\lambda}_k, \delta\hat{\lambda}_{k+n}\right) = \langle (\epsilon_{k+1} - \epsilon_k)(\epsilon_{k+n+1} - \epsilon_{k+n}) \rangle = D_\epsilon \times \begin{cases} 2, & n = 0, \\ (-1), & n = 1, \\ 0, & n \geq 2. \end{cases} \tag{119}$$

Thus, multiplicative measurement noise produces the same lag structure in $\hat{\mathcal{A}}_n(k)$ as additive independent measurement noise in (62): it is positive at lag $n = 0$, negative at lag $n = 1$, and vanishes for $n \geq 2$. The key difference is that the multiplicative contribution is independent of the cell age $k$, whereas for additive noise in the measured volume it decays with age through the factor $\langle v_k^{-2} \rangle$.

We now consider the accumulated growth. Using Eq. (115), we have

$$\hat{I}_k \equiv \log \frac{\hat{v}_k}{\hat{v}_b} = \log \frac{v_k\,(1+\epsilon_k)}{v_b\,(1+\epsilon_b)}. \tag{120}$$

Expanding to leading order in $\epsilon_k$, we obtain

$$\hat{I}_k = I_k + \delta\hat{I}_k + \mathcal{O}\left(\epsilon^2\right), \qquad \delta\hat{I}_k = \epsilon_k - \epsilon_b. \tag{121}$$

The contribution of multiplicative measurement noise to the accumulated growth autocovariance is therefore, using Eq. (121),

$$\delta\hat{\mathcal{B}}_n(k) \equiv \mathrm{Cov}\left(\delta\hat{I}_k, \delta\hat{I}_{k+n}\right) = \langle (\epsilon_k - \epsilon_b)\,(\epsilon_{k+n} - \epsilon_b) \rangle = D_\epsilon \times \begin{cases} 2, & n = 0, \\ 1, & n \geq 1. \end{cases} \tag{122}$$

Thus, multiplicative measurement noise also gives a particularly simple contribution to $\hat{\mathcal{B}}_n(k)$. For $n \geq 1$, it is a constant, just as in the additive iid case, but again without any dependence on the cell age $k$.

The main qualitative distinction between additive and multiplicative measurement noise is therefore their age dependence. For additive noise in the measured volume, the contribution to both $\hat{\mathcal{A}}_n(k)$ and $\hat{\mathcal{B}}_n(k)$ decreases with age because the same absolute error becomes less important as the cell grows. By contrast, multiplicative noise corresponds to a size-independent relative error, and hence its leading contribution is independent of $k$.

#### X. LINEAR GROWTH

In the main text, the growth rate is defined as

$$\lambda(t) = \frac{d \log v(t)}{dt}, \tag{123}$$

so that a constant growth rate corresponds to exponential growth in volume. An alternative description, relevant for linearly growing cells, is to define the growth rate directly from the time derivative of the volume,

$$g(t) \equiv \dot{v}(t). \tag{124}$$

In this case, the cell volume evolves as

$$v(t) = v_b + \int_0^t g(t')\,dt'. \tag{125}$$

The corresponding discrete-time growth observable is naturally defined by

$$g_k \equiv \frac{v_{k+1} - v_k}{\Delta t}. \tag{126}$$

For additive measurement noise

$$\hat{v}_k = v_k + \epsilon_k, \tag{127}$$

the measured growth rate is

$$\hat{g}_k \equiv \frac{\hat{v}_{k+1} - \hat{v}_k}{\Delta t} = g_k + \delta\hat{g}_k, \qquad \delta\hat{g}_k = \frac{\epsilon_{k+1} - \epsilon_k}{\Delta t}. \tag{128}$$

Thus, for additive measurement noise, the contribution of measurement noise has exactly the same form as in the multiplicative noise case discussed in Sec. IX.

Similarly to Eq. (25), we define the autocovariance

$$\hat{\mathcal{A}}_n^{\mathrm{lin}}(k) \equiv (\Delta t)^2 \operatorname{Cov}(\hat{g}_k, \hat{g}_{k+n}). \tag{129}$$

For independent additive measurement noise,

$$\langle \epsilon_k \rangle = 0, \qquad \langle \epsilon_k \epsilon_{k'} \rangle = D_\epsilon \delta_{k,k'}, \tag{130}$$

we obtain

$$\delta\hat{\mathcal{A}}_n^{\mathrm{lin}}(k) \equiv (\Delta t)^2 \operatorname{Cov}(\delta\hat{g}_k, \delta\hat{g}_{k+n}) = D_\epsilon \times \begin{cases} 2, & n = 0, \\ (-1), & n = 1, \\ 0, & n \geq 2. \end{cases} \tag{131}$$

Thus, for linear growth with additive measurement noise in the volume, the autocovariance of the instantaneous growth rate has the same lag structure as in the exponential growth case, but without the age-dependent prefactor $\langle v_k^{-2} \rangle$, compare with (50).

It is also natural to define the accumulated linear growth since birth by

$$J_k \equiv v_k - v_b = \int_0^{t_k} g(t')\, dt'. \tag{132}$$

The measured quantity is

$$\hat{J}_k = \hat{v}_k - \hat{v}_b = J_k + \delta\hat{J}_k, \qquad \delta\hat{J}_k = \epsilon_k - \epsilon_b. \tag{133}$$

We define

$$\hat{\mathcal{B}}_n^{\mathrm{lin}}(k) \equiv \operatorname{Cov}\left(\hat{J}_k, \hat{J}_{k+n}\right). \tag{134}$$

For independent additive measurement noise, we find

$$\delta\hat{\mathcal{B}}_n^{\mathrm{lin}}(k) \equiv \operatorname{Cov}\left(\delta\hat{J}_k, \delta\hat{J}_{k+n}\right) = D_\epsilon \times \begin{cases} 2, & n = 0, \\ 1, & n \geq 1. \end{cases} \tag{135}$$

Therefore, in the linear-growth setting with additive independent measurement noise, the contribution of measurement noise to both $\hat{\mathcal{A}}_n^{\mathrm{lin}}(k)$ and $\hat{\mathcal{B}}_n^{\mathrm{lin}}(k)$ is independent of the cell age $k$. This contrasts with the exponential-growth case with additive noise in the measured volume, where the same absolute error becomes less important as the cell grows.

More generally, if the additive measurement noise is temporally correlated,

$$\langle \epsilon_k \epsilon_{k+m} \rangle = C_m, \tag{136}$$

then the corresponding linear-growth formulas are

$$\delta\hat{\mathcal{A}}_n^{\mathrm{lin}}(k) = 2C_n - C_{n-1} - C_{n+1}, \tag{137}$$

$$\delta\hat{\mathcal{B}}_n^{\mathrm{lin}}(k) = C_n - C_k - C_{k+n} + C_0. \tag{138}$$

These expressions again show that linear growth leads to particularly simple measurement noise signatures, because no factors of the inverse cell volume appear.

#### XI. DETERMINISTIC TREND IN THE GROWTH RATE

In Secs. I and II, we considered an OU process with a constant mean growth rate. We now generalize the model by allowing a deterministic trend in the mean growth rate during the cell cycle.

Here, by a trend we mean a deterministic function of the cell age $t$, where $t$ is the time elapsed since birth. Thus, the trend is a function $\lambda_{\mathrm{trend}}(t)$, defined for $t \geq 0$, and the same function is used in every cell cycle. Importantly, this is not a trend that depends on the normalized age $t/\tau_j$, where $\tau_j$ is the division time of the $j$-th cycle. A normalized-age trend depends on the future division time of the same cycle, and is therefore conceptually different from the case considered here.

For a fixed lineage $L$, we consider the process

$$\dot{\lambda}(t) = -\gamma\left(\lambda(t) - \lambda_{\text{trend}}(t) - \ell_L\right) + \sqrt{2D_\lambda}\,\eta(t), \tag{139}$$

where $\ell_L \sim \mathcal{N}(0, D_\ell)$ is the lineage-dependent shift in the mean growth rate, as in Eq. (7), and $\eta(t)$ is Gaussian white noise with

$$\langle \eta(t) \rangle = 0, \qquad \langle \eta(t)\eta(t') \rangle = \delta(t - t'). \tag{140}$$

We define

$$\mu(t) \equiv \mathbb{E}\left[\lambda(t) | \ell_L\right] - \ell_L. \tag{141}$$

Thus, $\mu(t)$ is the expected growth rate within a given lineage after removing the constant lineage offset $\ell_L$. Since the noise has zero mean, taking the conditional expectation of Eq. (139) gives

$$\frac{d}{dt}\mathbb{E}\left[\lambda(t)|\ell_L\right] = -\gamma\left(\mathbb{E}\left[\lambda(t)|\ell_L\right] - \lambda_{\text{trend}}(t) - \ell_L\right), \tag{142}$$

and therefore

$$\dot{\mu}(t) = -\gamma\left(\mu(t) - \lambda_{\text{trend}}(t)\right). \tag{143}$$

Its solution is

$$\mu(t) = \mu(0)e^{-\gamma t} + \gamma \int_0^t e^{-\gamma(t-t')}\lambda_{\text{trend}}(t')\, dt'. \tag{144}$$

We now define the centered fluctuation

$$y(t) \equiv \lambda(t) - \mu(t) - \ell_L. \tag{145}$$

Substituting Eq. (145) into Eq. (139), and using Eq. (143), we find

$$\dot{y}(t) = -\gamma y(t) + \sqrt{2D_\lambda}\,\eta(t). \tag{146}$$

Thus, $y(t)$ obeys the same centered OU equation as in the case without a trend.

Its solution is

$$y(t) = y(0)e^{-\gamma t} + \sqrt{2D_\lambda}\int_0^t e^{-\gamma(t-t')}\eta(t')\, dt'. \tag{147}$$

Equivalently, the full process can be written as

$$\lambda(t) = \mu(t) + \ell_L + y(0)e^{-\gamma t} + \sqrt{2D_\lambda}\int_0^t e^{-\gamma(t-t')}\eta(t')\, dt'. \tag{148}$$

At division, the daughter cell is born with

$$\lambda^{(j+1)}(0) = \lambda^{(j)}(\tau_j) + \eta_d^{(j)}, \qquad \eta_d^{(j)} \sim \mathcal{N}(0, D_d), \tag{149}$$

as in Eq. (8). In terms of the centered fluctuation $y$, this gives

$$y^{(j+1)}(0) = y^{(j)}(\tau_j) + \mu(\tau_j) - \mu(0) + \eta_d^{(j)}. \tag{150}$$

Using Eq. (147) within the cycle at division time,

$$y^{(j)}(\tau_j) = y^{(j)}(0)e^{-\gamma\tau_j} + \sqrt{2D_\lambda}\,\zeta(\tau_j), \tag{151}$$

where $\zeta(\tau_j)$ is given in Eq. (13), we obtain

$$y^{(j+1)}(0) = y^{(j)}(0)e^{-\gamma\tau_j} + [\mu(\tau_j) - \mu(0)] + z_j, \qquad z_j = \sqrt{2D_\lambda}\,\zeta(\tau_j) + \eta_d^{(j)}. \tag{152}$$

Compared with the case without a trend, the only new term is the deterministic shift $\mu(\tau_j) - \mu(0)$. If, as in Sec. II, we replace the random cycle duration $\tau_j$ by its mean value $\langle\tau\rangle$, then this term becomes the constant $\mu(\langle\tau\rangle) - \mu(0)$. At stationarity, $\mu(0)$ must be chosen self-consistently so that $\mu(\langle\tau\rangle) = \mu(0)$. With this, the trend modifies the deterministic mean trajectory within the cycle, but does not affect the variance recursion. Therefore, under the same approximation used in Sec. II, the stationary birth variance remains

$$\mathrm{Var}\left(\lambda_b|\ell_L\right) = \mathrm{Var}\left(y_b|\ell_L\right) \approx \frac{D_\lambda}{\gamma} + \frac{D_d}{1 - e^{-2\gamma\langle\tau\rangle}}. \tag{153}$$

Accordingly, the autocovariance of the centered fluctuations is unchanged:

$$\mathrm{Cov}\left(y(t'), y(t'')|\ell_L\right) = \frac{D_\lambda}{\gamma}e^{-\gamma|t''-t'|} + \frac{D_d}{1 - e^{-2\gamma\langle\tau\rangle}}e^{-\gamma(t'+t'')}. \tag{154}$$

Averaging over lineages adds the same constant contribution $D_\ell$ as before, so the autocovariance of the full process is (compare with (24))

$$\mathrm{Cov}\left(\lambda(t'), \lambda(t'')\right) = \frac{D_\lambda}{\gamma}e^{-\gamma|t''-t'|} + \frac{D_d}{1 - e^{-2\gamma\langle\tau\rangle}}e^{-\gamma(t'+t'')} + D_\ell. \tag{155}$$

Therefore, within this approximation, introducing a deterministic trend $\lambda_{\text{trend}}(t)$ changes the mean growth rate trajectory during the cell cycle, but does not modify the autocovariance formulas used for $\mathcal{A}_n(k)$ and $\mathcal{B}_n(k)$ in Secs. III and IV. The effect of the trend is absorbed into the deterministic mean $\mu(t)$, while the fluctuations around that mean obey the same stochastic dynamics as before.

#### XII. DATA ANALYSIS FROM EXPERIMENTS

##### A. Preprocessing of the *E. coli* and mammalian data

For the *E. coli* data [3], we would like to discard filamentous cells. To this end, we discard cycles whose birth size $v_b$ or division size $v_d$ lies above the empirical 99.5% quantile of the corresponding empirical distribution. We note that the results reported in the main text are not affected if we keep the entire data.

For the mammalian data, we follow the authors of Ref. [4], and restrict the analysis to interphase and exclude mitosis from our analysis. Operationally, for each cycle we identify the onset of the $M$ phase and retain the cell cycle evolution only up to that boundary. The mass measurements in Ref. [4] are not taken at equal time intervals. However, our analysis requires equal time steps between measurements. We therefore interpolate each cell cycle onto an equally spaced $\Delta t'$ time grid. $\Delta t'$ is chosen to be the mean of all time intervals between measurements in the original data. Since this interpolation scheme introduces correlations between adjacent time points[1], we then omit every other time point, so we set $\Delta t = 2\Delta t'$. This yields equally spaced cell cycle trajectories that can be analyzed according to our framework.

##### B. Fitting the experimental data

From each retained cycle, we calculate the measured growth rate $\hat{\lambda}_k$ and the accumulated growth $\hat{I}_k$ according to Eqs. (42) and (51), respectively. From these quantities we compute the observables $\hat{\mathcal{A}}_n(k)$ and $\hat{\mathcal{B}}_n(k)$ defined in Eqs. (25) and (34).

---

[1] See Sec. VIII for a discussion about correlations in measurement noise.

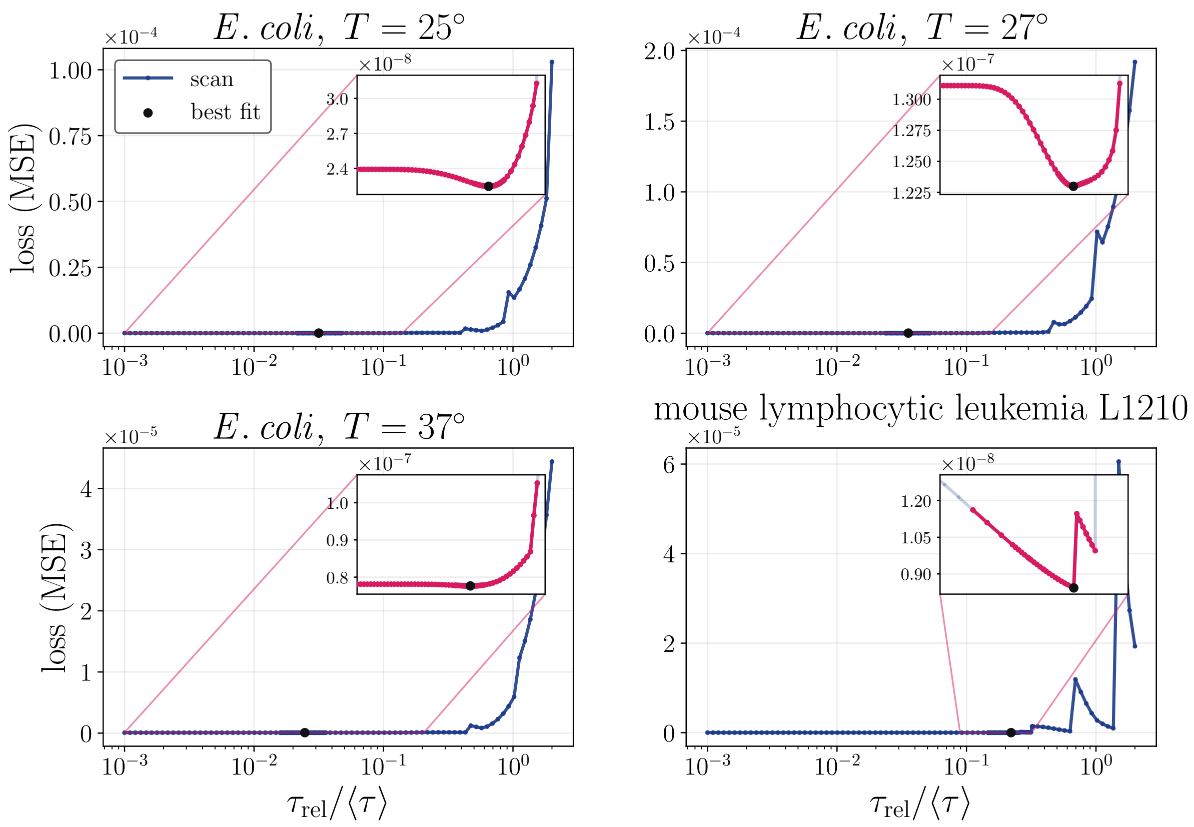


FIG. 1. The loss in Eq. (158) as function of $\tau_{\rm rel}/\langle\tau\rangle$, where $\tau_{\rm rel} = 1/\gamma$. For each value of $\tau_{\rm rel}$, the parameters $D_\lambda, D_d, D_\ell$ are fitted to the data $\hat{\mathcal{B}}_1^{\rm shift}$ to minimize the loss, and that minimal value is reported in this figure. The insets are a magnified view located around the minimal loss region.

As discussed in the main text, the data indicate that the measurement noise contribution dominates $\hat{\mathcal{A}}_0(k)$ at short times. We therefore estimate the measurement noise magnitude by fitting $\hat{\mathcal{A}}_0(k)$ to the exponential form given by Eq. (62). This fit yields an estimate of $D_\epsilon\langle v_b^{-2}\rangle$. We then define the corrected, "shifted", observable

$$\hat{\mathcal{B}}_1^{\rm shift}(k) = \hat{\mathcal{B}}_1(k) - D_\epsilon\langle v_b^{-2}\rangle, \tag{156}$$

which corresponds to the estimate of the biological contribution $\mathcal{B}_1 = \hat{\mathcal{B}}_1 - \delta\hat{\mathcal{B}}_1$.

We next fit $\hat{\mathcal{B}}_1^{\rm shift}(k)$ to $\mathcal{B}_1(k)$ within a fitting window determined by a survival threshold of 80%. For each fixed value of $\gamma$, we fit the remaining non-negative amplitudes $D_\lambda$, $D_d$, and $D_\ell$ in (recall Eqs. (38), (39), and (40))

$$\mathcal{B}_1(k) = \mathcal{B}_1^{\rm OU}(k;\gamma, D_\lambda) + \mathcal{B}_1^{\rm div}(k;\gamma, D_d) + \mathcal{B}_1^{\ell}(k; D_\ell). \tag{157}$$

The loss associated with a given $\gamma$ is the minimized mean squared residual over the fitting window,

$$L(\gamma) = \min_{D_\lambda, D_d, D_\ell \geq 0} \frac{1}{K_{\rm fit}} \sum_{k\in\mathcal{K}_{\rm fit}} \left[\hat{\mathcal{B}}_1^{\rm shift}(k) - \mathcal{B}_1(k)\right]^2, \tag{158}$$

where $\mathcal{K}_{\rm fit}$ denotes the selected fitting window and $K_{\rm fit}$ is the number of fitted time points. Equivalently, the scan can be parametrized by $\tau_{\rm rel}/\langle\tau\rangle = 1/(\gamma\langle\tau\rangle)$. The best fit corresponds to $\gamma$ that minimizes the loss in Eq. (158), see Fig. 1.

##### C. Bootstrap simulations

To estimate the uncertainty of the fitted parameters, we generate synthetic datasets using the best-fit values of $\gamma$, $D_\lambda$, $D_d$, $D_\ell$, and the inferred measurement noise magnitude $D_\epsilon$, and analyze them with exactly the same procedure as the experiment. The synthetic datasets are matched to the experimental sampling structure. In particular, we match the number of lineages and the number of cycles contributed by each lineage to those in the corresponding experiment.

For the *E. coli* data, the simulations use the inferred values of $\alpha$ and $\sigma_\xi$ (recall the size regulation model in Eq. (4)) using

$$\alpha = 1 - \mathrm{corr}(v_b, v_d) = 1 - \frac{\mathrm{Cov}\,(v_b, v_d)}{\sqrt{\mathrm{Var}(v_b)\mathrm{Var}(v_d)}}, \tag{159}$$

$$\sigma_\xi = 2\sqrt{\mathrm{Var}(v_b)\,\alpha(2-\alpha)}. \tag{160}$$

For the mammalian data, the simulations use the lineage structure of the experiment, the reported value of $\alpha = 0.55$ from [4], and a division noise amplitude $\sigma_\xi$ estimated from Eq. (160). In both cases, the simulated trajectories are sampled at the same effective time step as the data and are then processed in exactly the same way as the experimental trajectories.

Each synthetic realization is analyzed with the same fitting procedure as the experimental data. In particular, for every simulated dataset we re-estimate the measurement noise offset from $\hat{\mathcal{A}}_0(k)$, subtract this offset from $\hat{\mathcal{B}}_1(k)$, and only then perform the scan over $\gamma$ on the corrected observable $\hat{\mathcal{B}}_1^{\mathrm{shift}}(k)$ to minimize the loss (158). Thus, the bootstrap propagates uncertainty through the entire analysis procedure, including the measurement noise estimation step, rather than only through the final fit of $\mathcal{B}_1$.

##### D. Refitting the simulations and confidence intervals

For each fitted parameter, such as $\tau_{\mathrm{rel}}/\langle\tau\rangle$, $D_\lambda$, $D_d$, and, when relevant, $D_\ell$, the bootstrap yields an ensemble of refitted values. We summarize these ensembles by the histograms in Fig. 2. The black vertical line denotes the parameter value obtained from the experimental fit. The confidence intervals are taken from the empirical quantiles of the bootstrap ensemble, and we report the central 68% and 95% intervals.

In the mammalian data, the number of observed lineages is small. As a result, a fully parametric bootstrap in which the lineage-specific offsets are redrawn from a Gaussian $\mathcal{N}(0, D_\ell)$, with the fitted variance $D_\ell$, produces substantial realization-to-realization fluctuations. To avoid this source of variability, in the main analysis we use a conditional bootstrap.

In this conditional bootstrap, we first infer the lineage-specific mean growth rate offsets from the experimental data. We fit each cycle to an exponential, and then take the mean of the fitted growth rates within each lineage as the lineage-specific offset. We then keep these offsets fixed when generating the synthetic realizations. Thus, the bootstrap is conditioned on the observed lineage heterogeneity, rather than averaging over fresh draws from the fitted lineage-disorder distribution.

Each realization is nevertheless analyzed with the same fitting procedure used for the experimental data. Therefore, what is fixed is only the lineage-specific offset used in the simulation step. This procedure still allows us to estimate the uncertainty of the other parameters of the model, namely, $\gamma, D_\lambda, D_d$.

#### XIII. ADDITIONAL *E. COLI* DATASETS

Here we provide plots of the *E. coli* datasets taken from Ref. [3] for $T = 27°$, $37°$, see Figs. 3 and 4. We observe similar behavior and arrive at similar conclusions as for $T = 25°$ which is presented in the main text. See also Table I for the fitted values and dimensionless ratios $R_{\mathrm{birth}}, R_{\mathrm{cycle}}$ that measure the magnitude of division kicks with respect to the continuous OU noise.

TABLE I. Fitted parameters for all datasets. See Fig. 2. Confidence intervals are reported at the 95% level. The dimensionless ratio $R_{\mathrm{birth}} = D_d/(D_\lambda/\gamma)$ compares the division-kick variance to the OU growth rate variance at birth. The dimensionless ratio $R_{\mathrm{cycle}} = \frac{\mathcal{B}_0^{\mathrm{div}}(\langle\tau\rangle)}{\mathcal{B}_0^{\mathrm{OU}}(\langle\tau\rangle)} = \frac{D_d}{D_\lambda/\gamma}\frac{\left(1-e^{-\gamma\langle\tau\rangle}\right)^2}{2\left(\gamma\langle\tau\rangle-1+e^{-\gamma\langle\tau\rangle}\right)}$ compares the division kick and continuous OU contributions to accumulated growth fluctuations over one mean cell cycle. The ratios $R_{\mathrm{birth}}$ and $R_{\mathrm{cycle}}$ are computed from the best fit parameters.

| Dataset | Parameter | Fit Value | Confidence Interval | Unit |
|---|---|---|---|---|
| *E. coli* 25°C | $\tau_{\mathrm{rel}}/\langle\tau\rangle$ | 3.16 | [2.27, 5.01] | % |
| | $\tau_{\mathrm{rel}}$ | 2.10 | [1.33, 2.97] | min |
| | $\langle\tau\rangle$ | 66.6 | — | min |
| | $D_\lambda/\gamma$ | $9.68 \times 10^{-6}$ | $[3.75, 19.8] \times 10^{-6}$ | $\mathrm{min}^{-2}$ |
| | $D_d$ | $1.29 \times 10^{-4}$ | $[0.734, 2.83] \times 10^{-4}$ | $\mathrm{min}^{-2}$ |
| | $D_\ell$ | $5.65 \times 10^{-7}$ | $[1.94, 10.1] \times 10^{-7}$ | $\mathrm{min}^{-2}$ |
| | $R_{\mathrm{birth}}$ | 13.3 | — | – |
| | $R_{\mathrm{cycle}}$ | 0.218 | — | – |
| *E. coli* 27°C | $\tau_{\mathrm{rel}}/\langle\tau\rangle$ | 3.55 | [3.10, 6.82] | % |
| | $\tau_{\mathrm{rel}}$ | 1.86 | [1.43, 3.15] | min |
| | $\langle\tau\rangle$ | 52.4 | — | min |
| | $D_\lambda/\gamma$ | $2.41 \times 10^{-5}$ | $[0.956, 3.13] \times 10^{-5}$ | $\mathrm{min}^{-2}$ |
| | $D_d$ | $2.72 \times 10^{-4}$ | $[1.30, 4.79] \times 10^{-4}$ | $\mathrm{min}^{-2}$ |
| | $D_\ell$ | 0 | $[0, 7.20] \times 10^{-7}$ | $\mathrm{min}^{-2}$ |
| | $R_{\mathrm{birth}}$ | 11.3 | — | – |
| | $R_{\mathrm{cycle}}$ | 0.208 | — | – |
| *E. coli* 37°C | $\tau_{\mathrm{rel}}/\langle\tau\rangle$ | 2.47 | [1.61, 4.08] | % |
| | $\tau_{\mathrm{rel}}$ | 0.779 | [0.450, 1.14] | min |
| | $\langle\tau\rangle$ | 31.5 | — | min |
| | $D_\lambda/\gamma$ | $5.17 \times 10^{-5}$ | $[2.88, 10.7] \times 10^{-5}$ | $\mathrm{min}^{-2}$ |
| | $D_d$ | $7.08 \times 10^{-4}$ | $[3.91, 17.6] \times 10^{-4}$ | $\mathrm{min}^{-2}$ |
| | $D_\ell$ | $8.64 \times 10^{-7}$ | $[0.320, 14.9] \times 10^{-7}$ | $\mathrm{min}^{-2}$ |
| | $R_{\mathrm{birth}}$ | 13.7 | — | – |
| | $R_{\mathrm{cycle}}$ | 0.173 | — | – |
| Mouse lymphocytic leukemia L1210 | $\tau_{\mathrm{rel}}/\langle\tau\rangle$ | 22.1 | [10.9, 198] | % |
| | $\tau_{\mathrm{rel}}$ | 1.95 | [1.02, 18.9] | h |
| | $\langle\tau\rangle$ | 8.81 | — | h |
| | $D_\lambda/\gamma$ | $1.70 \times 10^{-4}$ | $[0, 3.33] \times 10^{-4}$ | $\mathrm{h}^{-2}$ |
| | $D_d$ | 0 | $[0, 2.96] \times 10^{-4}$ | $\mathrm{h}^{-2}$ |
| | $D_\ell$ | $9.94 \times 10^{-5}$ | $[0, 16.9] \times 10^{-5}$ | $\mathrm{h}^{-2}$ |
| | $R_{\mathrm{birth}}$ | 0 | — | – |
| | $R_{\mathrm{cycle}}$ | 0 | — | – |

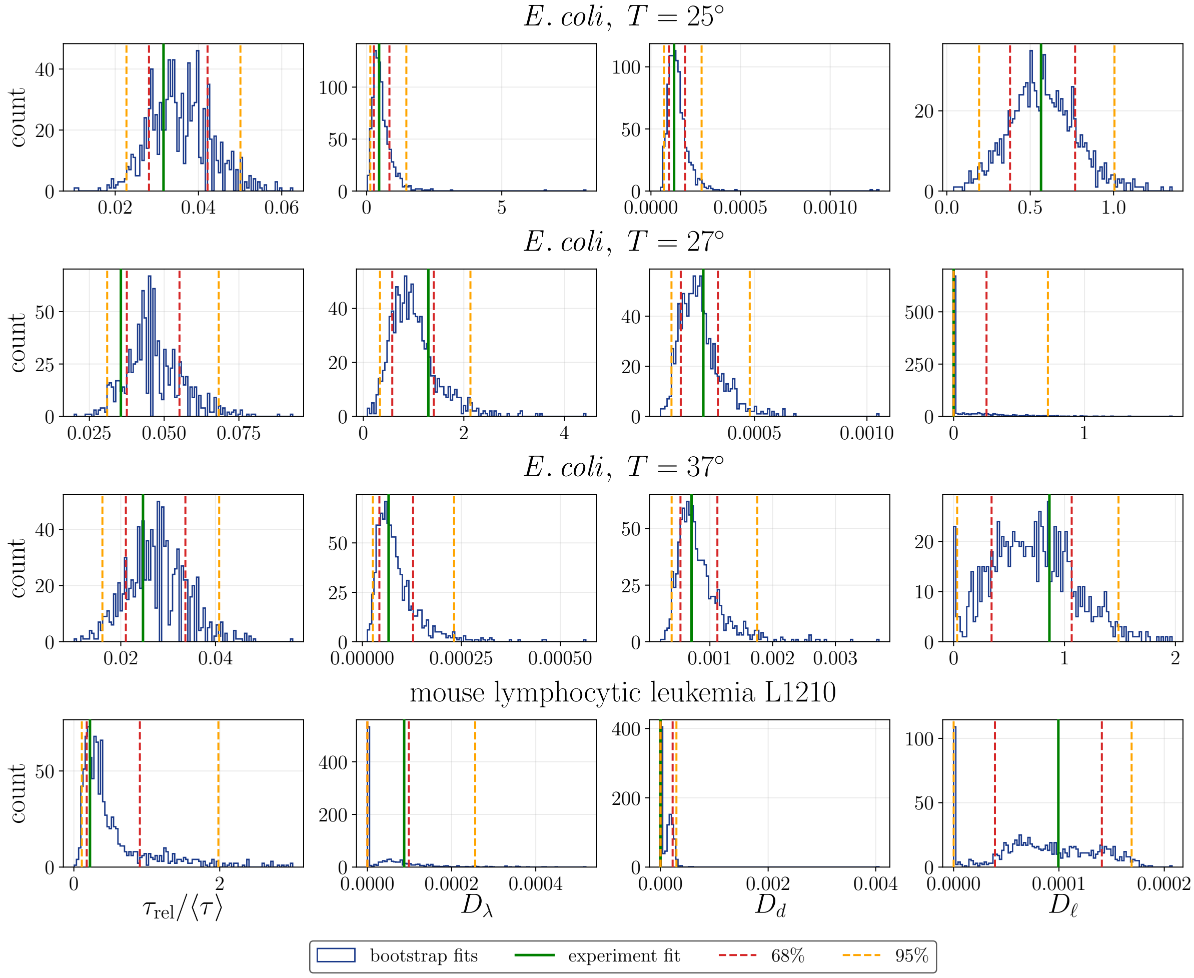


FIG. 2. Histograms of the fitted values from the simulations in Sec. XII C following the same procedure as performed on the experimental data in Sec. XII B. Given the fitted values from the experiments in Table I, denoted by a vertical green line in each panel, we perform 1000 simulated realizations. From the simulated data, we first estimate the noise magnitude from an exponential fit to $\hat{\mathcal{A}}_0$. Subsequently, we calculate $\hat{\mathcal{B}}_1^{\rm shift}$, fit it to $\mathcal{B}_1$, Eq. (157), by scanning a broad range of $\tau_{\rm rel} = 1/\gamma$, and reporting the parameters $D_\lambda, D_d, D_\ell$ which minimize the loss in Eq. (158). The blue histograms show the results of the fits for all simulations. The red and yellow lines mark the 68% and 95% quantiles.

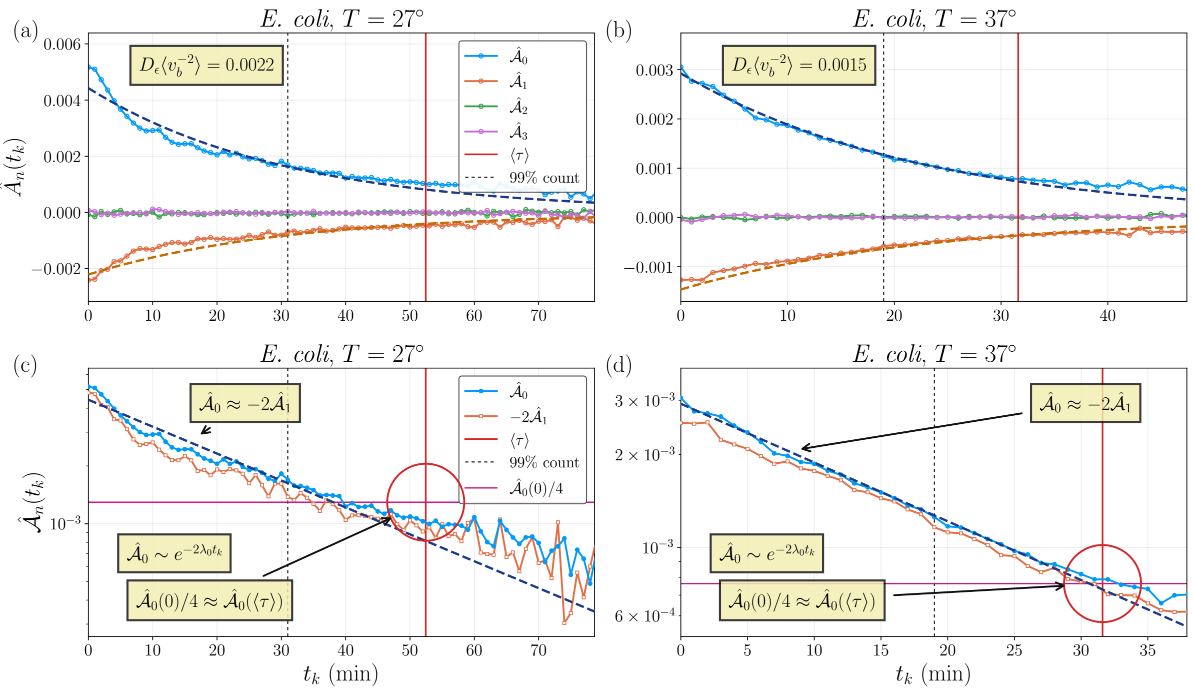


FIG. 3. Instantaneous growth rate autocovariance $\hat{\mathcal{A}}_n(k)$ for different lags $n$ for *E. coli* [3]. This Fig. corresponds to Fig. 2 in the main text for other temperature conditions from the same experiment. The top panels show that $\hat{\mathcal{A}}_n(k)$ follows the simple behavior predicted in Eq. (62): $\hat{\mathcal{A}}_0(k) > 0$, $\hat{\mathcal{A}}_1(k) < 0$, $\hat{\mathcal{A}}_{n\geq 2}(k) \approx 0$. The bottom panels show that $\hat{\mathcal{A}}_0(k) \approx -2\hat{\mathcal{A}}_1(k)$ (see Eq. (10) in the main text), $\hat{\mathcal{A}}_0(0)/4 \approx \hat{\mathcal{A}}_0(\langle\tau\rangle)$ (see Eq. (11) in the main text). Moreover, the dashed dark blue curves are exponential fits, $\hat{\mathcal{A}}_0(k) \approx 2D_\epsilon\langle v_b^{-2}\rangle e^{-2\lambda_0 t_k}$, which provide an estimate for the measurement noise magnitude. The orange dashed line in the top panel corresponds to $-D_\epsilon\langle v_b^{-2}\rangle e^{-2\lambda_0 t_k}$, using the parameters from the fit of $\hat{\mathcal{A}}_0(k)$. For *E. coli* at $T = 27°$ we find $D_\epsilon\langle v_b^{-2}\rangle \approx 0.0022$ and for $T = 37°$ we find $D_\epsilon\langle v_b^{-2}\rangle \approx 0.0015$ .

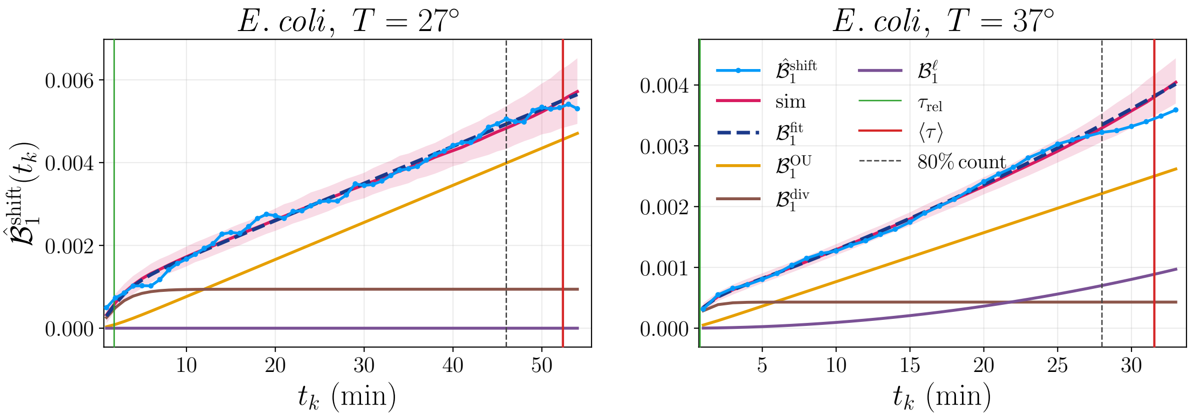


FIG. 4. The shifted accumulated growth rate autocovariance $\hat{\mathcal{B}}_1^{\text{shift}}(k) \equiv \hat{\mathcal{B}}_1(k) - D_\epsilon\langle v_b^{-2}\rangle$ for *E. coli* [3]. This Fig. corresponds to Fig. 3 in the main text for other temperature conditions from the same experiment. The constant $D_\epsilon\langle v_b^{-2}\rangle$ is obtained from the fit in Fig. 3. The blue points are the data, and the dark blue dashed lines are fits of $\hat{\mathcal{B}}_1^{\text{shift}}$ using Eq. (158) performed as described in Sec. XII B. From the fit, we plot separately the different contributions: OU noise (yellow), division kicks (brown), and lineage variability (purple), see legend in panel (a). In pink, we plot the median of 1000 realizations of simulations performed with the values of $\gamma, D_\lambda, D_d, D_\epsilon, D_\ell$ obtained from the experiments, as described in Secs. XII C, XII D. The shaded pink ribbon marks the 95% confidence interval. The green line denotes the fitted relaxation time $\tau_{\text{rel}} = 1/\gamma$. The red line is the mean division time $\langle\tau\rangle$. The vertical black dashed line marks the threshold for the data we used for the fit. We set the threshold to 80% count of the initial number of cells in order to mitigate the effects of survival bias.

[1] Ariel Amir. Cell Size Regulation in Bacteria. *Physical Review Letters*, 112(20):208102, May 2014.

[2] Po-Yi Ho, Jie Lin, and Ariel Amir. Modeling Cell Size Regulation: From Single-Cell-Level Statistics to Molecular Mechanisms and Population-Level Effects. *Annual Review of Biophysics*, 47(1):251–271, May 2018.

[3] Yu Tanouchi, Anand Pai, Heungwon Park, Shuqiang Huang, Nicolas E. Buchler, and Lingchong You. Long-term growth data of Escherichia coli at a single-cell level. *Scientific Data*, 4(1):170036, March 2017.

[4] Ethan Levien, Joon Ho Kang, Kuheli Biswas, Scott R. Manalis, Ariel Amir, and Teemu P. Miettinen. Stochasticity in mammalian cell growth rates drives cell-to-cell variability independently of cell size and divisions. *Proceedings of the National Academy of Sciences*, 123(10):e2516372123, March 2026.
